# Characterisation of RNA editing and gene therapy with a compact CRISPR-Cas13 in the retina

**DOI:** 10.1101/2024.02.10.579778

**Authors:** Satheesh Kumar, Yi-Wen Hsiao, Vickie H Y Wong, Deborah Aubin, Jiang-Hui Wang, Leszek Lisowski, Elizabeth P Rakoczy, Fan Li, Luis Alarcon-Martinez, Anai Gonzalez-Cordero, Bang V Bui, Guei-Sheung Liu

## Abstract

CRISPR-Cas13 nucleases are programmable RNA-targeting effectors that can silence gene expression in a reversible manner. Recent iterations of Cas13 nucleases are compact for adeno-associated virus (AAV) delivery to achieve strong and persistent expression in various organs in a safe manner. Here, we report significant transcriptomic signatures of Cas13bt3 expression in retinal cells and show all-in-one AAV gene therapy with Cas13bt3 can effectively silence *VEGFA* mRNA in human retinal organoids and humanised *VEGF* transgenic mouse (trVEGF029, Kimba) models. Specifically, human embryonic stem cells (hESC)-derived retinal pigment epithelium cells show high expression of Cas13bt3 from virus delivery corresponding to a significant reduction of *VEGFA* mRNA. We further show that intravitreal delivery of Cas13bt3 can transduce mouse retinal cells efficiently, reaching the photoreceptors for specific knockdown of human *VEGFA* in the Kimba mouse. Our results reveal important considerations for assessing Cas13 activity and establish Cas13bt3 as a potential anti-VEGF agent that can achieve long-term control of VEGFA for the treatment of retinal neovascularization.

## Introduction

The emergence of Clustered Regularly Interspaced Short Palindromic Repeats (CRISPR) and CRISPR-associated (Cas) enzymes, known as CRISPR-Cas, has fuelled several gene editing efforts for novel therapeutics in the past decade(1). Developed by repurposing the bacterial immune defence against phage viruses, CRISPR-Cas harnesses the ability to induce targeted changes to DNA or RNA sequences. Among CRISPR-Cas proteins, CRISPR-Cas13 is of particular interest for its exclusive specificity for RNA. Targeting RNA is especially attractive as it enables control of gene expression without perturbing the genome and introducing permanent changes and is therefore an emerging avenue for therapeutic development(2).

Eleven variants of CRISPR-Cas13 have been described to date(3). Among them, Cas13d (commonly referred to as CasRx) described in 2018, garnered special attention for its compact size of 930aa, making it the first Cas13 enzyme amenable to adeno-associated virus (AAV) delivery. Studies investigating the development of RNA-targeting therapeutics have widely used CasRx to demonstrate preclinical efficacy against disorders of vision, hearing and brain function(4–7), however significant unintended editing has been reported, hampering its clinical progress(8, 9), and the exact specificity profile of CasRx remains contested(10). More recently, hypercompact Cas13bt3 (also reported as Cas13X) enzymes were described, ranging even smaller at 775aa(11, 12). The newfound enzyme has been reported to possess high specificity, and its small size enables further applications such as RNA base editing using a single AAV(13–15). The therapeutic potential of Cas13bt3 RNA silencing is yet to be demonstrated, however initial *in vitro* studies show promising efficiency and safety(13, 16).

As a leading cause of blindness in the elderly population affecting 200 million people globally, ocular neovascularization refers to a group of conditions characterised by the abnormal growth of leaky blood vessels in the eye. Vascular endothelial growth factor (VEGF) has been described as a key mediator of this process, and the overexpression of *VEGFA* underlies pathological angiogenesis and neovascularization in common ocular diseases like diabetic retinopathy (DR) and neovascular age-related macular degeneration (nAMD). Regular (monthly/bimonthly) administration of anti-VEGF drugs is the approved front-line treatment modality for these patients(17). Delivered through intravitreal injections, frequent administration of anti-VEGF drugs poses a significant clinical burden on healthcare institutions and patients due to the risk of infection, patient anxiety, clinic capacity and availability of trained healthcare personnel among others(18).

We have previously described a potential alternative therapy to current anti-VEGF injections using CRISPR-CasRx gene therapy for the treatment of ocular neovascularization. This approach was validated *in vitro* in HEK293FT and MIO-M1 cells, demonstrating more than 90% and 50% *VEGFA* mRNA knockdown, respectively(19). Here, we used Cas13bt3 to demonstrate potent *VEGFA* knockdown *in vitro* in human cell lines, 3D human embryonic stem cells (hESC)-derived retinal organoids, and *in vivo* in transgenic humanised mouse retinas. Promisingly, our results show a significant knockdown of *VEGFA* mRNA in human cell lines without impact on predicted off-target genes. Single-cell RNA sequencing (scRNA-seq) was used to characterise Cas13bt3-mediated *VEGFA* mRNA knockdown at single cell resolution in human retinal organoids identifying, for the first time, the overall transcriptomic impact in retinal cells. Lastly, *in vivo* gene therapy showed robust *VEGFA* mRNA silencing in mouse models of proliferative retinopathy along with control of neovascularization symptoms. Overall, these findings identify key considerations for developing CRISPR-Cas13 ocular gene therapies and demonstrate a potential clinical application as an anti-VEGF agent, potentially offering an alternative to current treatment options.

## Results

### CRISPR-Cas13bt3 can knock down VEGFA mRNA in mammalian cells with minimal unintended targeting activity

We used our previously described single guide RNAs (sgRNA) targeting Exon 4 of *VEGFA* mRNA (19) to develop the Cas13bt3 plasmids to assess RNA cutting efficiency followed by collateral and off-target cleavage (**Figure 1A-D**). Collateral activity is the *trans*-cleavage of nearby mRNA transcripts by Cas13 enzymes after activation (**Figure 1B**). Off-target activity on the other hand, is the targeting of non-target mRNA transcripts due to unspecific (or partially specific) binding of the sgRNA (**Figure 1C**).

**Figure 1.**
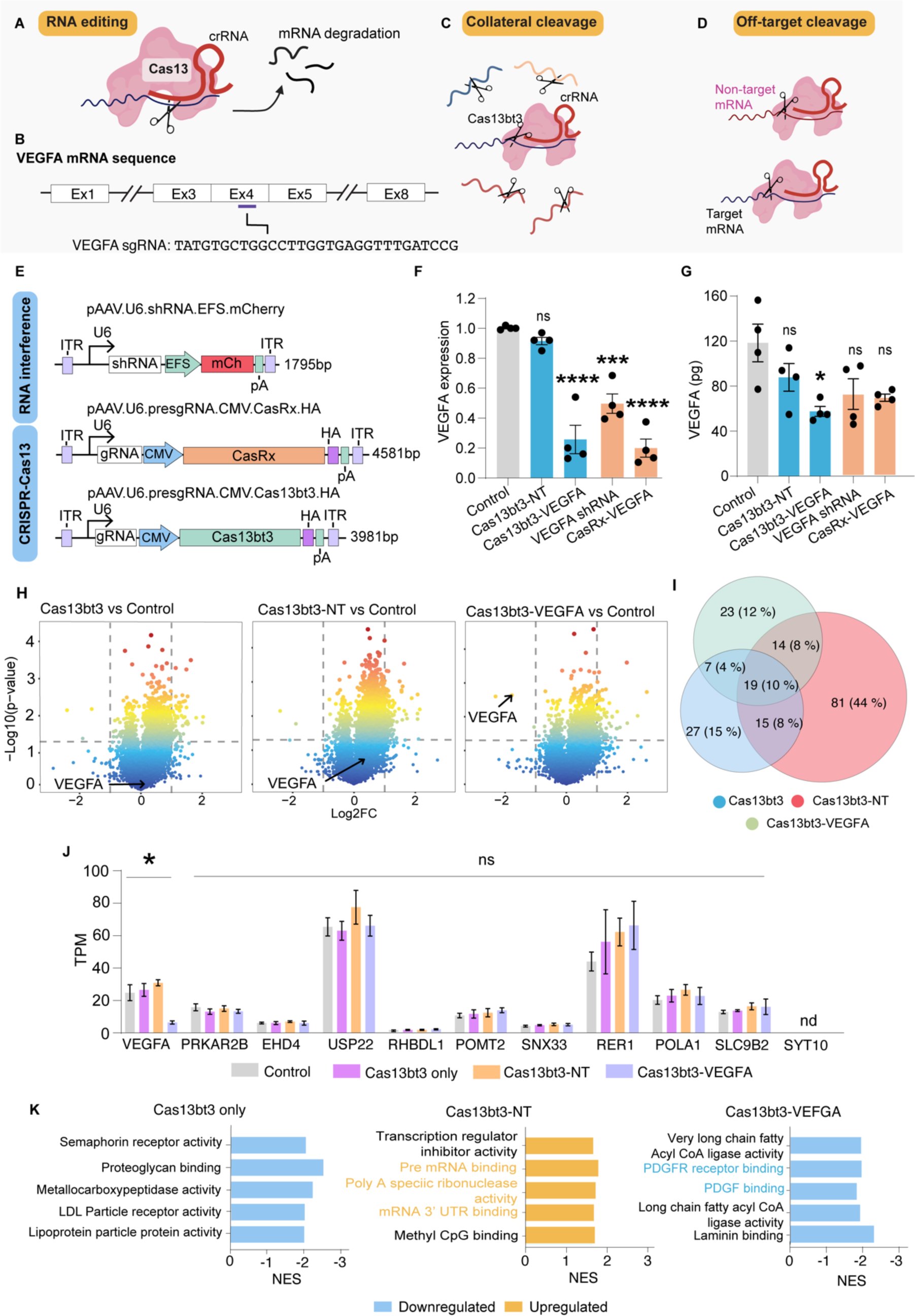
Comparison of RNA-targeting tools for in vitro testing of VEGFA mRNA silencing. **(A)** Schematic of CRISPR-Cas13 mechanism of action. **(B)** VEGFA sgRNA sequence and targeting region on VEGFA mRNA sequence. **(C)** Schematic of collateral and **(D)** off-target cleavage. **(E)** Schematic for domain structure of VEGFA targeting shRNA and Cas13 (CasRx and Cas13bt3) vectors. **(F)** VEGFA mRNA knockdown from shRNA, CasRx and Cas13bt3 in HEK293FT cells (n = 4). (G) VEGFA protein expression from ELISA after transfection of shRNA, CasRx and Cas13bt3 plasmids (n = 4). **(H)** Volcano plots showing DEGs from Cas13bt3 only, Cas13bt3-NT sgRNA and Cas13bt3-VEGFA sgRNA compared to transfection control in HEK293FT cells. VEGFA gene is denoted by a black arrow. **(I)** Venn diagram of a total number of differentially expressed genes (DEGs) from Cas13bt3 only, Cas13bt3-NT sgRNA and Cas13bt3-VEGFA sgRNA plasmids. **(J)** TPM plot for VEGFA and candidate off-target genes. **(K)** Top 5 enriched GO molecular function (MF) terms from Cas13bt3 only, Cas13bt3-NT sgRNA and Cas13bt3 VEGFA sgRNA. NES above zero represents significant activation of the pathway, while NES below zero represents significant suppression of the pathway. Orange text denotes key factors that are upregulated; blue text denotes key factors that are downregulated. Data presented as mean ± SEM. Statistical analysis was conducted by one-way ANOVA and Tukey’s multiple comparison test **(F, G)** and Welch’s t-test **(J)**. ns: not significant, *p≤0.05, **p≤0.01, ***p≤0.001, ****p≤0.0001.

We sought to compare and validate the efficiency of the Cas13bt3 enzyme by testing RNA cutting against two previously established RNA-targeting methods, RNA interference (short hairpin RNA, shRNA) and CasRx (a variant of Cas13d). The final all-in-one AAV plasmids carrying either *VEGFA*-targeting shRNA or precursor single guide RNA (sgRNA hereafter) with either CasRx or Cas13bt3 (**Figure 1D**) were transfected into HEK293FT cells. Here, to further enhance the expression of Cas13bt3, we incorporated the stronger cytomegalovirus (CMV) promoter instead of the previously used elongation factor 1α short (EFS) promoter (19).

Significant *VEGFA* mRNA knockdown was observed from all three RNA-targeting methods, with up to 70% shRNA *VEGFA* mRNA knockdown (*p* < 0.001), while CasRx and Cas13bt3 achieved up to 90% knockdown (*p* < 0.0001) compared to vector controls (AAV plasmid carrying a fluorescent marker) (**Figure 1F**; n = 4 independent experiments). Next, we sought to determine if the reduction in mRNA would also lead to a decrease in protein levels. Interestingly, ELISA for VEGFA revealed that only the Cas13bt3 treatment led to a significant reduction of VEGFA protein in the cell supernatant (**Figure 1G**, *p* < 0.05; n = 4 independent experiments). While it is unclear why only the Cas13bt3-VEGFA sgRNA group showed a significant reduction in protein levels, it is well-established that mRNA transcripts are not predictive of the corresponding protein levels (20). Interestingly, cell lysate samples revealed that VEGFA protein could not be detected across all groups, indicating that most VEGFA molecules were exported outside the cells.

To identify the collateral activity of Cas13bt3 on the transcriptome, bulk RNA sequencing (RNA-seq) was performed after HEK293FT cells were treated with either vector control, Cas13bt3 only, Cas13bt3-NT or Cas13bt3-VEGFA constructs. Volcano plots of Cas13bt3 samples against control demonstrated and confirmed significant downregulation of *VEGFA* mRNA only in cells that received the Cas13bt3-VEGFA treatment (**Figure 1H).** Interestingly, we noted the highest number of differentially expressed genes (DEGs) from the Cas13bt3-NT sgRNA group compared to Cas13bt3 only or Cas13bt3-VEGFA (81 vs 27 and 23, respectively; **Figure 1H**), a finding that corresponds with recent studies (16, 21). Overall, the DEG data suggested minimal collateral activity from the Cas13bt3-VEGFA comparable to Cas13bt3 only. Moreover, we also explored the collateral activity of Cas13bt3 compared to shRNA and CasRx. A similar level of DEGs was found in all 3 groups (**Figure S1A-E**), indicating that collateral activity from the Cas13bt3 is comparable to previously established RNA-targeting methods.

To understand off-target activity from Cas13, genes with high complementary regions to the targeting sgRNA need to be identified. We therefore identified potential off-target genes using NCBI BLAST for the VEGFA sgRNA sequence and studied the expression levels of the top 10 candidate off-target genes (**Figure S2A**). Promisingly, none of the off-target genes were significantly downregulated, indicating a lack of off-target activity from Cas13bt3 (**Figure 1J**), as was the case for shRNA and CasRx (**Figure S1F**).

To further examine the DEG levels between the NT and targeting sgRNAs, we identified the top significantly enriched gene ontology (GO) terms for the Cas13bt3-NT sgRNA and Cas13bt3-VEGFA sgRNA groups. Top 5 significant GO terms for the Cas13bt3-NT sgRNA group were all positive enrichments and specifically showed upregulation of RNA binding processes, such as “*pre-mRNA binding*” and “*mRNA 3’ UTR binding*”, possibly explaining the elevated levels of DEGs in this group. On the other hand, with the VEGFA sgRNA, the top 5 GO terms were all negative enrichments and particularly showed downregulation of “*platelet-derived growth factor (PDGF) binding*” and “*PDGF receptor binding*”, both angiogenesis promoting factors (**Figure 1K**). A recent study showed CasRx to have intrinsic targets that affected essential processes like RNA metabolism and mitotic cell cycle (21) and opined that this could underlie the lethality seen upon CasRx expression in a mouse model (22). Here, we found that laminin-related processes (e.g., “*laminin binding*”, “*laminin complex*”) were strongly downregulated across all groups (Cas13bt3 only, Cas13bt3-NT sgRNA and Cas13bt3-VEGFA sgRNA) suggesting intrinsic targets could exist for Cas13bt3 as well (**Data S1**). While we did not observe significant toxicity in our cell cultures, whether this is detrimental to cellular health and overall host organism needs to be investigated.

### Multiplexing sgRNAs for CRISPR-Cas13bt3 knockdown of the target gene from a single-AAV vector

The small size of Cas13bt3 and the significant downregulation of *VEGFA* mRNA observed prompted us to hypothesize that multiple sgRNAs can be delivered using a single AAV vector to increase overall targeting and RNA knockdown efficiency. In theory, multiple sgRNAs would increase the rate of Cas13 activation and levels of intracellular Cas13-sgRNA complexes for RNA silencing.

To understand Cas13bt3 activation and demonstrate multiplex targeting, we transfected Cas13bt3-VEGFA sgRNA and Cas13bt3 with a VEGFA sgRNA array (carrying three VEGFA-targeting sgRNAs) (**Figure 2A-C**). Here, cells transfected with Cas13bt3-VEGFA sgRNA or Cas13bt3-VEGFA sgRNA array showed similar levels of significant *VEGFA* mRNA knockdown (<90%) (**Figure 2D**; n = 3 independent experiments), indicating no improved knockdown from the multiple sgRNA system. We then sought to study if delivering multiple sgRNAs will increase overall transcriptomic impact and performed RNA-seq to similarly determine collateral and off-target cleavage. Overall, DEGs for both single and multiple sgRNA groups were comparable and suggested minimal collateral activity from the sgRNA array system compared to the Cas13bt3-NT group (**Figure 2E**). Again, we identified top candidate off-target genes for the other *VEGFA* sgRNAs (**Figure S2B and C**) and did not detect any significant off-target activity in these genes by the Cas13bt3-VEGFA sgRNA array system as well (**Figure 2F**). Previously, with CasRx, we observed moderately lower knockdown rates with a sgRNA array (19). With Cas13bt3 however, multiplexing sgRNAs did not seem to increase or dampen editing efficiency. We speculate active Cas13 nucleases competing for target mRNA or low levels of *VEGFA* mRNA transcripts may have resulted in the lack of improvement in targeting efficiency. The additional sgRNAs may also have poor targeting capabilities resulting in no overall net effect. Still, our results suggest that multiple sgRNAs can be used simultaneously with Cas13bt3 while avoiding significant collateral activity.

**Figure 2.**
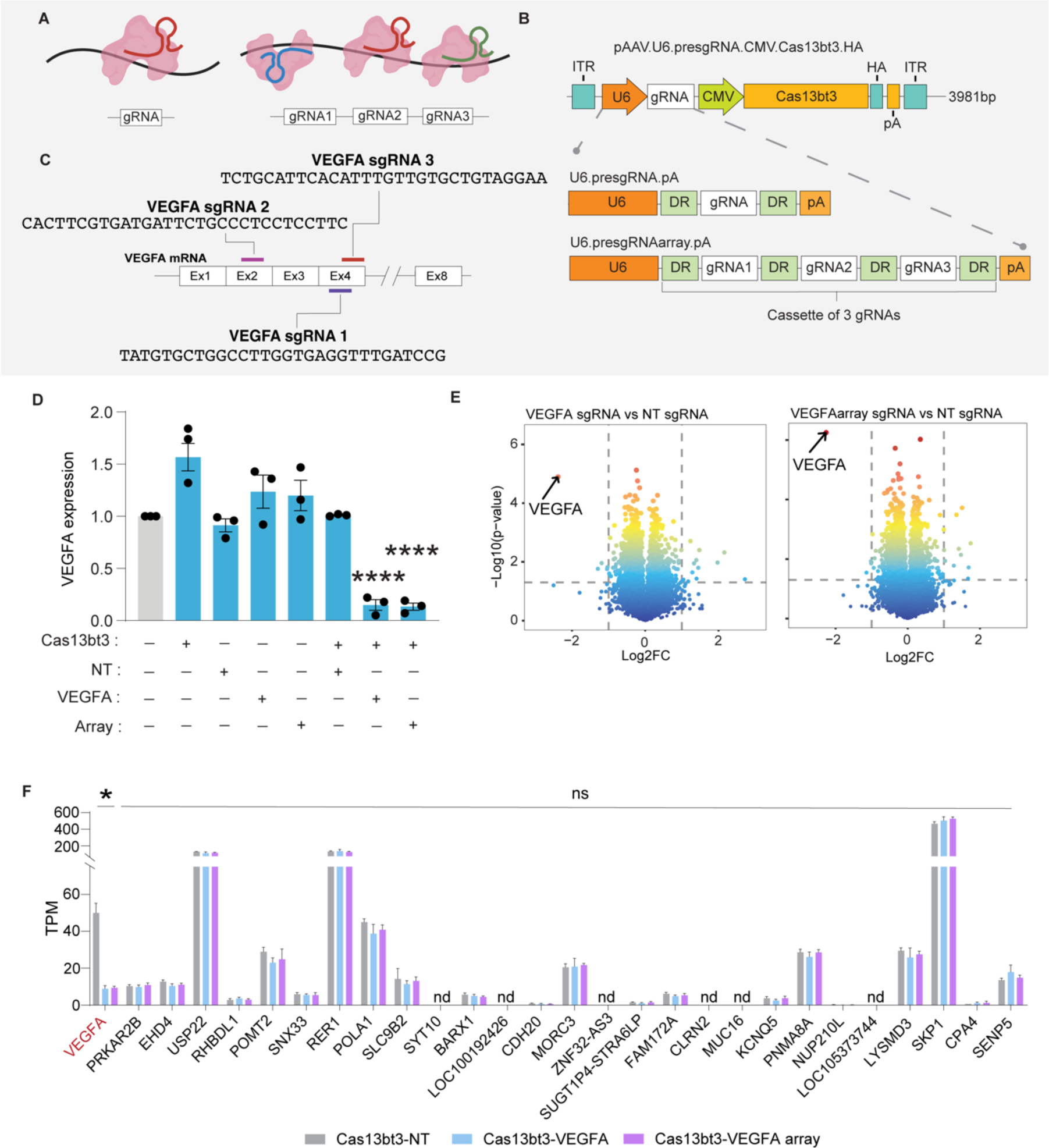
Guide-dependent and guide-independent off-targets from Cas13bt3-mediated VEGFA knockdown. **(A)** Schematic of single and multiplex targeting with Cas13bt3 and **(B)** U6-sgRNA domain in single sgRNA and multiple sgRNA vectors. **(C)** Guide RNA targeting regions in VEGFA mRNA. **(D)** VEGFA mRNA knockdown from Cas13bt3, non-targeting and targeting sgRNAs. **(E)** Volcano plots showing differentially expressed genes between Cas13bt3 with non-targeting and **(F)** TPM plots for VEGFA and candidate off-target genes. Target gene VEGFA is highlighted in red. Data presented as mean ± SEM. Statistical analysis was conducted by one-way ANOVA and Tukey’s multiple comparison test **(D)** and Welch’s t-test **(F)**. ns: not significant, *p≤0.05, ***p≤0.001, ****p≤0.0001.

Apart from competition for substrate mRNA, it was recently reported that the presence of “UC” sites on the target mRNA may also affect knockdown efficiency, with “UC” sites within 10-20 bases of spacer binding to allow for optimal cleavage (**Figure S3A**)(16). We found that while sgRNA1 and sgRNA2 had “UC” sites in these optimal regions, sgRNA3 had only one distal “UC” site 40 nucleotides away from the spacer binding region (**Figure S3B**). While we did not independently test the knockdown efficiency of our sgRNAs, this finding suggests that sgRNA 3 alone would achieve low levels of RNA knockdown. As no improved knockdown was observed from the VEGFA sgRNA array, we proceeded to investigate Cas13bt3 with a single sgRNA further.

### AAV2.7m8 delivery efficiently expresses Cas13bt3 in various retinal cells

Human organoids are excellent models for gaining clinically relevant information on new therapeutic agents, as they are derived from human pluripotent stem cells (PSCs) and reproduce morphological, functional and transcriptomic features *in vitro*(23). Recently, hESC-derived retinal organoids have been used to investigate eye development and model inherited retinal diseases, as well as the testing of novel gene therapies(24, 25). While CRISPR-Cas9 has entered the clinic, CRISPR-Cas13-based therapies are still being extensively explored at the preclinical stage, and demonstrating the utility of Cas13 in retinal organoids will be a major advance for ocular gene therapy.

Here, we used our well-established 2D/3D differentiation protocol to generate hESC-derived retinal organoids containing all the major cell types in the retina (**Figure S4**)(26). First, to ensure maximum expression of the Cas13bt3 in the retinal cells, we codon optimised the Cas13bt3 protein sequence to obtain two other variants of Cas13bt3 (Cas13bt3_COP1_ and Cas13bt3_COP2_) (**Figure S5A**). We compared the expression levels of the original Cas13bt3 with Cas13bt3_COP1_ and Cas13bt3_COP2_ by transfection in HEK293FT cells and found both variants to exhibit increased expression and knockdown efficiency (**Figure S5B-D**). Among them, Cas13bt3_COP1_ (Cas13bt3 hereafter) showed the highest expression and knockdown and was used for subsequent investigation. To elucidate the activity of Cas13bt3 and efficacy for AAV gene therapy in *bona fide* human retinal cells, we transduced 19 weeks old organoids with AAV2.7m8.U6.NTsgRNA.Cas13bt3 and AAV2.7m8.U6.VEGFAsgRNA.Cas13bt3 viral vectors. Retinal organoids were then harvested for single cell RNA sequencing (scRNA-Seq) 6 weeks after treatment (∼25 weeks; **Figure 3A**). First, we confirmed that the major retinal cell types were clustered similarly across experimental conditions (S1: Untreated; S2: AAV2.7m8.U6.VEGFAsgRNA.Cas13bt3; and S3: AAV2.7m8.U6.NTsgRNA.Cas13bt3) (**Figures 3B and S6**, n = 3 pooled organoids per condition). We used our recently described retinal atlas dataset to determine the fidelity of retinal organoids and identified the expression of the different retinal cell markers(27), confirming exclusive cell-specific expression of the markers (**Figure S7**).

**Figure 3.**
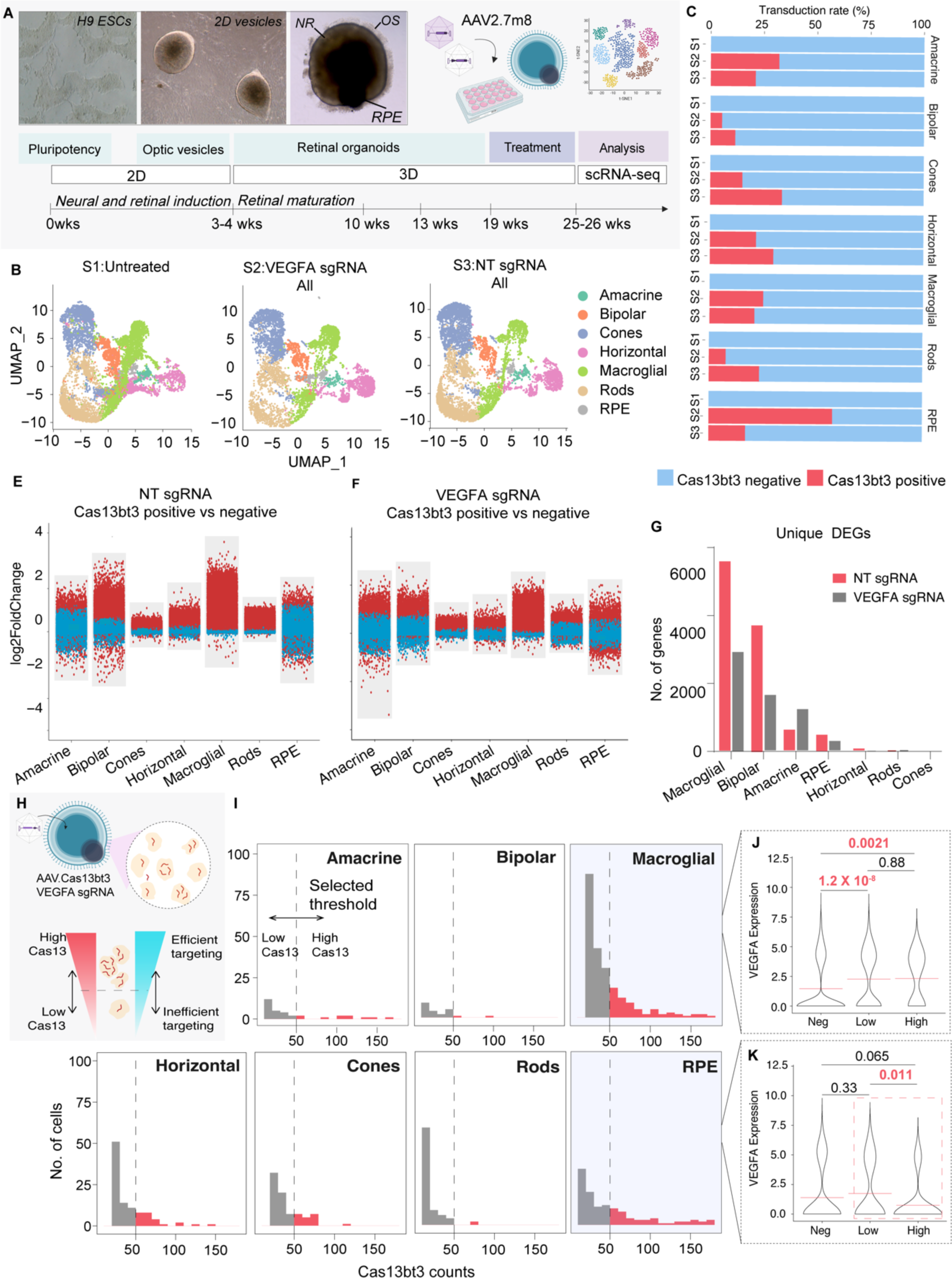
Cas13bt3 treatment and single-cell RNA sequencing of 3D retinal organoids. (A) Schematic of 3D retinal organoid scRNA-seq experimental procedure. (B) UMAP plots show retinal cell type clusters for the untreated, NT sgRNA and VEGFA sgRNA treatment groups. (D) Infection rate of retinal organoids by cell type and treatment group. (E) Multi-volcano plots showing differentially expressed genes (DEGs) between Cas13bt3 positive and negative cells from the NT sgRNA and (F) VEGFA sgRNA treatment groups. (G) Number of unique DEGs across retinal cell types. **(H)** Schematic of variable Cas13bt3 expression in retinal organoid cells and relationship between CRISPR-Cas13bt3 expression and targeting efficiency. **(I)** Histograms showing the level of Cas13bt3 expression by number of reads across retinal cell types in the VEGFA sgRNA group. **(J)** Violin plots of VEGFA expression against the level of Cas13bt3 expression for macroglial cells and **(K)** RPE cells. Red box indicates cells of interest for high *VEGFA* targeting activity.

Next, to examine Cas13bt3 expression in the retinal organoids, we evaluated read numbers and classified cells containing more than 10 reads of Cas13bt3 to be positive for viral infection. Using this threshold, we determined an average transduction efficiency of 18.78% (1403 out of 7467 cells) for the VEGFA sgRNA group and 25.95% (2260 out of 8706 cells) for the NT sgRNA group. Next, we analysed the transduction and the presence of Cas13bt3 in specific cell types. While all cell types showed noticeable levels of Cas13bt3, RPE cells showed high levels of expression with over 50% of cells containing Cas13bt3 in the VEGFA sgRNA (S2) treatment group (153 out of 264 cells). This is likely due to the reported preferential transduction of RPE cells by the AAV2.7m8 serotype (28). Interestingly, in the NT sgRNA group, only ∼15% of RPE cells expressed Cas13bt3, which could owe to varying culture conditions. Bipolar cells showed the least expression of Cas13bt3 (≤10%), while all other cell types showed moderate expression (≤25%) (**Figures 3C and S8**). Overall, our results indicated AAV2.7m8 as a useful delivery vector for cells across retinal cell types, confirming several previous studies (29–32).

To investigate the impact of the Cas13bt3 expression in retinal cells, we compared DEGs between cells positive and negative for Cas13bt3 in the retinal organoid that received the AAV2.7m8.U6.NTsgRNA.Cas13bt3 or AAV2.7m8.U6.VEGFsgRNA.Cas13bt3. Here, we observed a significant number of DEGs across all cell types in the retinal organoid that received the AAV2.7m8.U6.NTsgRNA.Cas13bt3, where most DEGs were upregulated genes (**Figure 3E**). The same pattern was observed in the retinal organoid transduced with AAV2.7m8.U6.VEGFAsgRNA.Cas13bt3 (**Figure 3F**), albeit at a lower extent, suggesting that the NT sgRNA exhibited a greater impact on the transcriptome. To determine the specific cell types that were more sensitive to Cas13bt3 expression, we identified unique DEGs from each cell type (to remove common effects from AAV delivery and culturing conditions) across the retinal cells and found macroglial and bipolar cells to exhibit the highest DEGs, which was especially higher in the NT sgRNA group (Macroglial - NT: 5571 vs VEGFA: 2928, Bipolar – NT: 3725 vs VEGFA: 1633) (**Figure 3G**). These results emphasised the need to consider the effect of non-targeting sgRNAs when assessing Cas13 specificity.

### Cas13bt3 can significantly knock down VEGFA mRNA in retinal organoids

Next, we examined Cas13bt3 targeting efficiency and its effect on *VEGFA* expression across the retinal cell types. First, we determined *VEGFA* expression in control untreated organoids. While Cas13bt3 positive cells have been identified, the abundance of Cas13bt3 reads within cells may still vary (e.g. 10 reads vs 100 reads) and affect the determination of targeting efficiency. We hypothesized that increased intracellular Cas13bt3 transcripts allow for higher targeting efficiency and thereby increased silencing of *VEGFA* mRNA (**Figure 3H**). We first examined the number of Cas13bt3 reads in cells that received AAV2.7m8.U6.NTsgRNA.Cas13bt3 and AAV2.7m8.U6.VEGFAsgRNA.Cas13bt3 treatments (**Figures 3I and S9**). To accurately determine the Cas13bt3 effect and to understand the abundance of the targeting complex within each cell, we examined AAV2.7m8.U6.VEGFAsgRNA.Cas13bt3 treated cells. We selected a threshold to classify high (≥ 50 reads) and low (<50 reads) expression of Cas13bt3 that enables evaluation of targeting efficiency. We found that RPE and macroglial cells expressed high levels of Cas13bt3 expression, while most cones, rods, and horizontal and bipolar cells only showed low levels of Cas13bt3 expression (**Figure 3I**).

We then quantified *VEGFA* mRNA expression in cells expressing high and low levels of Cas13bt3. To this end, we pooled cells within the high and low Cas13bt3 thresholds and compared average *VEGFA* expression against Cas13bt3 negative cells to determine if loss of *VEGFA* is dependent on high levels of Cas13bt3. A significant decrease of *VEGFA* in high Cas13bt3-expressing cells compared to low Cas13bt3-expressing cells would indicate VEGFA silencing from increased targeting activity, while a significant reduction compared to negative cells would indicate *VEGFA* loss from viral delivery. A prominent observation was that viral delivery appeared to upregulate *VEGFA* expression in macroglial cells with both low and high Cas13bt3-expressing cells exhibiting a significant increase in VEGFA expression (**Figure 3J**). Moreover, we observed that RPE cells were the only cells to show a significant reduction of *VEGFA* expression with high Cas13bt3 expression compared to low Cas13bt3 cells (*P* < 0.05) (**Figures 3K**, red square, **and S10**). When we correlated Cas13bt3 reads to VEGFA expression, again, only RPE cells that received the AAV2.7m8.U6.VEGFAsgRNA.Cas13bt3 treatment showed a significant reduction (*p* = 0.043) (**Figure S11**).

### *VEGFA* knockdown impedes disease progression in a humanised mouse model of proliferative retinopathy

Lastly, we sought to investigate if *VEGFA* knockdown can control neovascularization *in vivo* in mouse models of proliferative retinopathy. This offers a potential new tool against eye diseases causing neovascularization. To this end, we utilised the trVEGF029 (*VEGFA*^+/-^, Kimba) mouse, a humanised transgenic mouse model where the neuroretina overexpresses h*VEGFA* leading to neovascularization like human proliferative DR(33). To ensure that optimal transduction of the retina is being achieved, we delivered a mCherry reporter gene under the control of CMV promoter using the AAV2.7m8 serotype, known to have excellent transduction of the retina when delivered intravitreally into C57BL/6 mice (**Figure S12A and B**). Immunohistochemistry analysis of AAV2.7m8.CMV.mCherry treated retinal flatmounts showed high levels of fluorescence through all retinal layers (RGCs, bipolar and photoreceptor cells) 4 weeks following injections, confirming quick onset and deep penetration of the retina from intravitreal delivery (**Figure S12C**; n = 6 eyes from 3 animals).

Next we treated Kimba mice with AAV2.7m8 viruses driving our codon optimised Cas13bt3 carrying NT sgRNA (AAV2.7m8.U6.NTsgRNA.Cas13bt3) or VEGFA sgRNA (AAV2.7m8.U6.VEGFAsgRNA.Cas13bt3). Six-week-old Kimba mice received bilateral, intravitreal injections of AAV2.7m8.U6.NTsgRNA.Cas13bt3 or AAV2.7m8.U6.VEGFAsgRNA.Cas13bt3. Eight weeks following treatment, the eyes were analysed for *VEGFA* mRNA expression, neovascular growth, and retinal degeneration (**Figure 4A**). Similar to the control mCherry treatments, high transduction of photoreceptor cells was observed as demonstrated by co-localisation of mCherry and HA-tag (Cas13bt3) with Rhodopsin positive cells in the ONL (**Figure 4B and C**). In addition, qPCR analysis showed high expression of Cas13bt3 in Kimba retinas injected with AAV.2.7m8.U6.NTsgRNA.Cas13bt3 and AAV2.7m8.U6.VEGFAsgRNA.Cas13bt3 (**Figure 4D**; n = 3-12 retinas). This corresponded with ∼50% reduction (*p* < 0.001) of human *VEGFA* (*hVEGFA)* mRNA (**Figure 4E**), but not mouse *VEGFA* (*mVEGFA*) (**Figure 4F**) in the AAV2.7m8.U6.VEGFAsgRNA.Cas13bt3 group, while no reduction of *hVEGFA* or *mVEGFA* was observed with AAV2.7m8.U6.NTsgRNA.Cas13bt3 or no treatment (**Figure 4E and F**; n = 3-12 retinas). These results confirmed the specific knockdown of *hVEGFA* from Cas13bt3 despite the high complementarity of human and mouse *VEGFA* sequences (29 out of 30 nucleotides) in the targeting region (**Figure 4G**). RNA sequencing further confirmed that *mVEGFA* expression and predicted off-target genes were not significantly altered, indicating low off-target activity from Cas13bt3-mediated RNA silencing (**Figure S13**).

**Figure 4.**
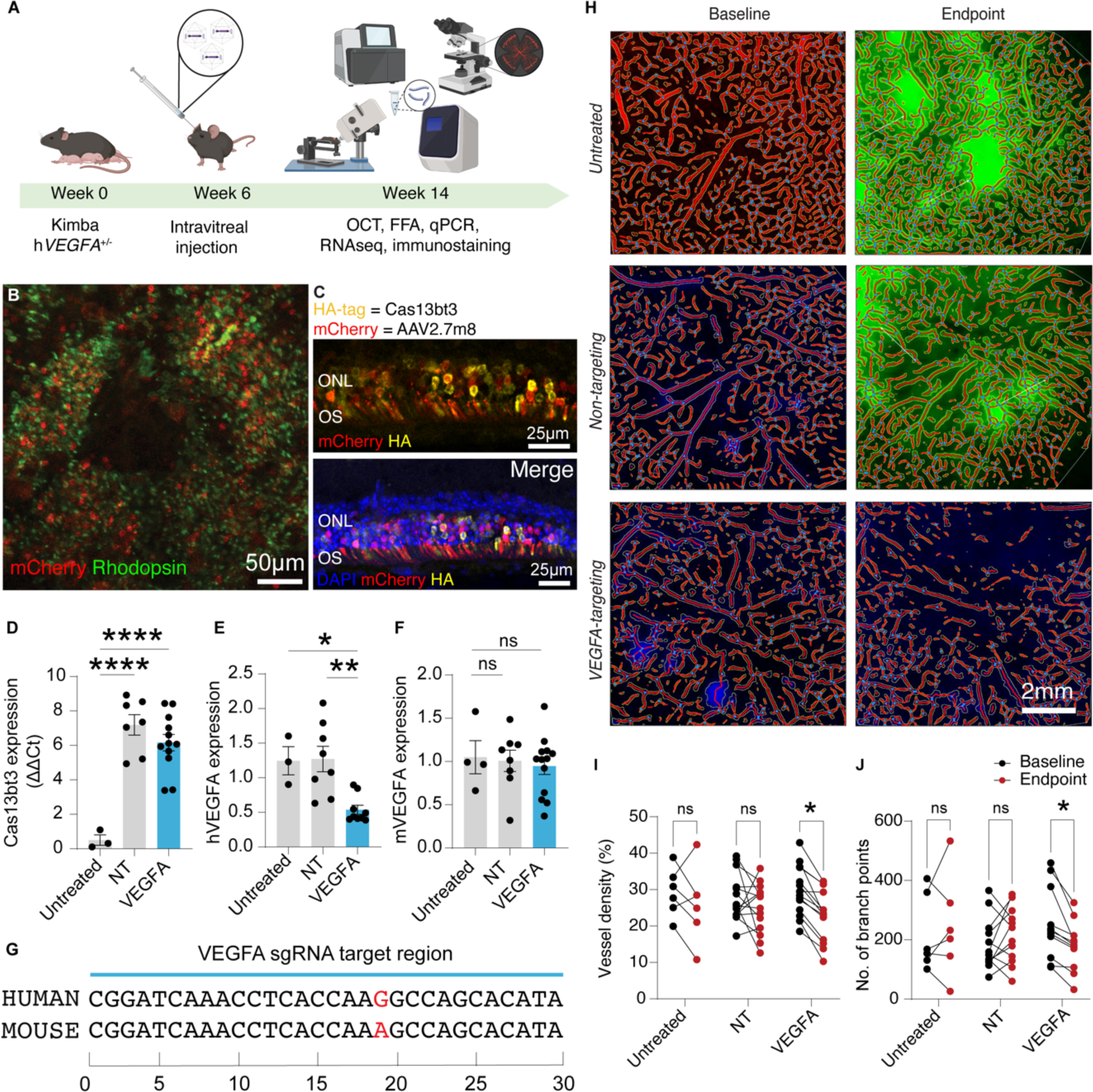
*In vivo* VEGFA mRNA knockdown in humanized Kimba mice. **(A)** Schematic of experimental procedure for intravitreal treatment of Kimba mice. **(B and C)** Immunohistochemistry of retinal flatmount and cryosection showing transduction of AAV2.7m8 and expression of Cas13bt3 (Magnification: 20X and 40X, respectively). **(D)** Expression of Cas13bt3 from Kimba mice retinas. **(E)** Expression of human and **(F)** mouse VEGFA mRNA in Kimba retinas. **(G)** VEGFA mRNA targeting regions in human and mouse transcriptome. **(H)** Central fluorescein fundus angiography (FFA) images obtained from Kimba mice at baseline (6 weeks) and endpoint (14 weeks) after AngioTool analysis and quantification. **(I)** A plot of vessel density and **(J)** number of branch points from untreated and treated Kimba eyes at baseline (6 weeks) and endpoint (14 weeks) was analyzed using in vivo central fluorescein fundus angiography (FFA). Images were edited using Adobe Photoshop to remove central retinal arteries and veins. Green spots and white arrows indicate leakage from blood vessels. Data presented as mean ± SEM. Statistical analysis was conducted by one-way ANOVA with Tukey’s multiple comparison test **(D-F)** and Multiple paired t-tests **(I-J)**. *p≤0.05, **p ≤0.01, ****p≤0.0001.

To confirm if *VEGFA* mRNA knockdown influenced disease progression, we analysed fundus fluorescein angiography (FFA) images from Kimba eyes for vessel leakage and quantified vessel density and number of branch points, which are signs indicative of the neovascular process. We used fluorescein to label vessels of the AAV2.7m8.U6.VEGFAsgRNA.Cas13bt3, AAV2.7m8.U6.NTsgRNA.Cas13bt3-treated and untreated eyes. We observed large spots of vessel leakage, indicating neovascularization had progressed in both untreated and NTsgRNA-treated eyes (**Figures 4H and S14**; green expression and white arrows). However, in the AAV2.7m8.U6.VEGFAsgRNA.Cas13bt3 group, no leakage was observed (**Figures 4H and S14**; VEGFA targeting). Immunohistochemistry against laminin-2 was used to image through the retinal vascular layers (superficial, intermediate, and deep vascular plexuses) (**Figure S15A and B**). Blood vessels appeared diminished in the intermediate and deep vascular layers for the VEGFA sgRNA group. In the NT sgRNA group, the neovascular process could be observed through leaky vessel structures in the same layers (**Figure S15C**; n = 4 retinas). As disease progression in each retina is variable, we performed paired analyses of recordings from each retina at the endpoint to its baseline. The AAV2.7m8.U6.VEGFAsgRNA.Cas13bt3 treated eyes showed significantly lower vessel density and branch points compared to baseline at 6 weeks post-treatment, unlike the untreated and the AAV27m8.U6.NTsgRNA.Cas13bt3 eyes (**Figure 4I and J**; n = 4-13 retinas, *p* < 0.05). Next, to determine whether *VEGFA* knockdown leads to rescue of overall retinal structure. we performed optical coherence tomography (OCT) to analyse total retinal thickness (TRT) before and after treatment. Unexpectedly, we observed no significant changes between the untreated and treated groups and a general decline in TRT across all animals (**Figure S16**). We postulate that the overexpression of *hVEGFA* promotes rapid retinal degeneration that could not be controlled or reversed through *VEGFA* silencing.

Overall, our findings showed that Cas13bt3-mediated RNA silencing significantly reduces VEGFA mRNA expression, specifically targeting *hVEGFA* and controlling hallmark signs of neovascularization in a humanised mouse model. This approach promises a useful alternative to current anti-VEGF drugs for proliferative retinopathy.

## Discussion

Our comprehensive *in vitro*, *ex vivo* and *in vivo* characterisation of Cas13bt3 activity and performance in the retina provides novel insights for the therapeutic use of the enzyme. Cas13bt3 was first described only two years ago but has garnered significant attention for its compact size and reported specificity(11). Structural characterization and activation mechanisms have recently been described, which has spurred immense interest in the molecule for therapeutic development(16, 34). Due to ongoing controversy over the exact specificity of Cas13 nucleases(10), we aimed to characterise Cas13bt3 efficiency and specificity in mouse and human cells. We chose to target the *VEGFA* gene as this growth factor is essential for both physiological and pathological angiogenesis. In the retina, physiological functions include its role as a survival factor for retinal ganglion cells(35), photoreceptors, Müller cells(36), and RPE(37). In disease, overexpression of *VEGFA* leads to neovascularization and vision loss. Therefore, the clinical applicability of *VEGFA* downregulation in specific cell types of the retina is of great importance. Previous studies have demonstrated transcript knockdown using CasRx proteins(11, 12). Notably, CasRx-mediated *VEGFA* mRNA knockdown has also been shown to reduce choroidal neovascularization after silencing *VEGFA* mRNA in RPE cells *in vivo* in a laser-induced choroidal neovascularization model(4). Our results confirmed that Cas13bt3 mediates significant knockdown of *VEGFA* in human cell lines, hESC-derived retinal cells, and the mouse retina.

*In vitro*, and specifically in HEK293FT cells, several studies have shown Cas13 nucleases to be highly precise(10). Most of these studies, however, compared differentially expressed genes to a NT sgRNA control, not accounting for transcriptomic perturbations from the Cas13-NT sgRNA complex. A recent study showed high collateral activity from CasRx when compared to an empty vector(38). Here, we compared targeting Cas13bt3 to vector controls showing minimal, but positive collateral activity. More importantly, we show that Cas13bt3 with a NT sgRNA has a significant transcriptomic impact, resulting in high levels of DEGs. While Cas13bt3 may undergo activation after maturation of a NT sgRNA; without adequate homologous pairing, the catalytic pocket cannot be formed for efficient nuclease activity (39). On the other hand, with a targeting gRNA, Cas13bt3 would be activated to form the Cas13bt3-gRNA-target complex for targeted knockdown. Our results suggest that with a NT sgRNA, Cas13bt3 initiates an unspecific target-search process involving significant RNA binding that greatly impacts the expression levels of several genes, while a targeting sgRNA narrows the target-search process to reduce overall transcriptomic impact. We observed that despite high levels of overall DEGs, the number of downregulated genes remained similar between the NT sgRNA and VEGFA sgRNA groups suggesting that the NT sgRNA does not induce the formation of a catalytic pocket for RNA knockdown and altered gene expression is likely a result of increased RNA binding. Comparison of different NT sgRNAs may further elucidate this phenomenon. Overall, these findings suggest that NT sgRNAs are not appropriate controls for determining the specificity of Cas13 nucleases, raising awareness against such evaluation.

Multiplexed RNA targeting with Cas13bt3 was also demonstrated by incorporating multiple sgRNAs within a single AAV vector for targeted knockdown of *VEGFA* mRNA. While this appears to be an attractive approach to enhance transcript knockdown, our study established that multiple sgRNAs failed to improve transcript degradation. The rate of mRNA knockdown was comparable to single sgRNA treatment. Previously, we similarly observed when investigating multiplexed RNA editing with CasRx carrying three *VEGFA*-targeting sgRNAs, the multiple sgRNA system unexpectedly led to decreased knockdown efficiency(19), raising the suspicion that substrate competition or inadequate Cas13 activation plays an inhibitory role in RNA targeting efficiency. Interestingly, the comparable levels of DEGs in both single and multiple sgRNA vectors suggest that Cas13bt3 activation is not drastically elevated with the presence of additional sgRNAs. While we did not investigate the delivery of sgRNAs targeting multiple genes, our results show multiple sgRNAs do not increase overall collateral activity, which might prove useful when several genes must be targeted simultaneously to control disease progression. This approach could potentially be applied even to ocular neovascularization by simultaneously targeting the *VEGFA* and *PDGF* genes for enhanced treatment(40).

For the first time, we also investigated AAV-mediated Cas13 RNA editing in retinal organoids at single cell resolution. While several studies have demonstrated *VEGFA* silencing for control of neovascularization, previous studies have focused on specific cell types that are believed to be major contributors to *VEGFA* expression, like RPE and Müller cells(4, 41). While this proves useful to demonstrate proof-of-concept, it does not provide a comprehensive understanding relevant to therapeutic development as *VEGFA* silencing in one cell type may have collateral effects on other retinal cells. Retinal organoid models enable the testing of therapies in an environment that mimics the human retina containing lamination and major retinal cell types of the developing retina. Here we sought to understand the retina-wide effect following the delivery of Cas13bt3 as well as the targeted silencing of *VEGFA*. Our findings not only demonstrate the use of Cas13bt3 for ocular gene therapy but also bring to light several considerations for its use. First, we showed transduction across all retinal cell types studied with the clinically relevant AAV2.7m8 vector confirming its effectiveness as a vector for retinal transduction and supporting its clinical utility(28, 31). Among the retinal cell types studied, bipolar cells showed the least transduction by AAV2.7m8, while RPE cells seemed more susceptible. It is likely that these differences are due to cell-specific transduction as RPE cells are readily transduced by a variety of AAV capsids, particularly AAV2.7m8(29). It should also be noted that overall transduction was determined using an arbitrary cut-off (Cas13bt3 read counts ≥ 10) based on our previous experience with the retinal organoids(26).

Like our *in vitro* findings, NT sgRNA activity following gene therapy led to numerous DEGs when compared to VEGFA targeting sgRNA. This emphasises the need for a well-designed targeting sgRNA for reduced collateral activity. Retinal organoids showed significantly more DEGs than HEK293FT cells, which is expected partly due to cellular response to virus delivery. Still, notwithstanding response to AAV, the higher DEG levels also indicate retinal cells to be more sensitive to Cas13, demonstrating the importance of studying nuclease activity and efficiency in physiologically relevant models, an important consideration for CRISPR-Cas-mediated gene therapies. As preclinical studies with Cas13 nucleases are commonly performed in HEK293FT cells, its specificity has often been exaggerated in the literature. Our results strongly suggest that Cas13 specificity must be evaluated in relation to untreated or delivery vector controls in organ-specific cells to accurately determine safety.

On the front of therapeutic potential, we could only confirm significant targeted *VEGFA* mRNA knockdown in RPE cells *ex vivo*, likely due to preferential transduction of AAV2.7m8 and the consequent strong expression of Cas13bt3 in these cells. Lower copies of Cas13bt3 in the other cells may have resulted in the lack of significant *VEGFA* knockdown, and further studies are required to understand the levels of Cas13 expression necessary in other retinal cells for effective activity. Testing of cell type-specific promoters could also be beneficial as the CMV promoter is known for its susceptibility to silencing in certain cell types, such as stage-specific PSCs-derived photoreceptor cells (42). Importantly, with RPE cells being the main producers of *VEGFA* in nAMD, our results show that Cas13bt3 gene therapy could be a potential anti-VEGF tool for other types of neovascular eye diseases.

Lastly, we tested the anti-VEGF potential of Cas13bt3 in the retina of a *VEGF* transgenic mouse model of proliferative retinopathy, the Kimba mouse. We opted to examine the anti-VEGF potential of Cas13bt3 in Kimba mice as it shows chronic disease and all major signs of proliferative retinopathy (33). Importantly, as Kimba mice express human *VEGFA*, the same vectors tested in retinal organoids could also be used *in vivo*. Following RNA editing gene therapy in Kimba retinas, we demonstrated the specific knockdown of human *VEGFA*, with no significant impact on endogenous mouse *VEGFA*. This shows strong evidence for the specificity of targeted Cas13bt3 activity. Silencing of *VEGFA* mRNA also rescued vascular phenotypes showing an improvement in controlling vessel growth and leakiness. The major concern with current anti-VEGF treatment is the need for frequent intravitreal injections. Recent efforts to address this problem have employed gene editing strategies like targeting *VEGFA* using CRISPR-Cas9 (41, 43). However, the efficacy of such treatments is limited by the efficient and safe delivery of vectors due to the size of Cas9, the poor efficiency of compact Cas9 nucleases, and the risk of genomic alterations from the long-term expression of Cas9(44). Our results indicate that a single intravitreal injection could achieve durability and potency while reversible silencing of *VEGFA*, addressing drawbacks with current gene editing strategies. While RNA editing is generally considered safer than DNA-targeted options, it would be prudent to consider the effects of long-term Cas13 expression, and Cas13 systems amenable to regulation are of consideration for therapeutic development.

There are a few limitations to our study. First, we did not extensively investigate the stress responses we observed from the retinal cells. While HEK293FT appears resistant to high levels of collateral activity, retinal cells exhibited significantly higher levels of transcriptomic effects. This is supported by the high levels of DEGs we observed in various cells from the retinal organoids, suggesting significant inflammatory responses as well as cell-specific collateral activity. We also found the upregulation of ubiquitin-like protein (UBL) processes across the cell types in all groups regardless of VEGFA silencing, strongly indicating increased cellular stress from delivery of Cas13. As RNA-targeting molecules like Cas13 require persistent expression for therapeutic effect, this could pose a significant safety concern and must be further studied. This could be done through studying the expression of essential growth factors and developmental and functional genes after treatment. Second, while vessel density and branching were noticeably reduced, we could not extensively study the recovery of phenotype in our mouse model due to the severity of the disease. OCT readings showed significant retinal degeneration at the treatment timepoint, making rescue of phenotype challenging to assess. Lastly, we were also unable to confirm the reduction of VEGFA protein from the mice retinas as not enough protein could be obtained for quantitative analysis.

CRISPR-Cas technology has advanced at breakneck speed in the last decade, revolutionising medicine and offering potential treatment opportunities for numerous previously uncurable diseases. The recent FDA approval of an *ex vivo* CRISPR therapy for sickle-cell disease (branded Casgevy) is evidence of this progress. However, a comprehensive understanding of the CRISPR nucleases is crucial. In this study, we offer novel insights into the Cas13 targeting mechanism and its utility in targeting retinal and RPE cells, providing evidence for the potential of Cas13-based therapies for ocular disease. This study demonstrated that the hypercompact Cas13bt3 can be efficiently delivered and expressed in various retinal cell types for effectively controlling *VEGFA* expression and neovascularization. An in-depth evaluation of collateral activity in retinal cells would facilitate the development of a safe and effective Cas13-mediated anti-VEGF AAV gene therapy.

## Materials and Methods

### Plasmid construction

The protein sequences of CasRx and Cas13bt3 were gifts from P. Hsu(45) and H. Yang(11), respectively. Cas13bt3 was codon optimized using online tools from Integrated DNA Technologies (IDT), GenScript and Benchling. The all-in-one AAV plasmids for shRNA, CasRx and Cas13bt3 were designed as previously described(19), and synthesized by GenScript. All plasmids used in this study are available from Addgene.

### Cell culture and transfections

HEK293FT cells (American Type Culture Collection, ATCC) were maintained in Dulbecco’s Modified Eagle’s Medium (DMEM; Thermo Fisher Scientific) supplemented with 10% (v/v) fetal bovine serum (FBS, Bovogen Biologicals) and 1% (v/v) antibiotic-antimycotic (Thermo Fisher Scientific) in a humidified 5% CO_2_ incubator at 37°C.

### RT-qPCR

At 48 hours after transfection, RNA was extracted from cells using the Monarch^®^ Total RNA Miniprep Kit (New England Biolabs and 200ng of RNA was converted to complementary DNA (cDNA) using High-Capacity cDNA Reverse Transcription Kit (Thermo Fisher Scientific) according to the manufacturers’ instructions. cDNA was diluted with water at the ratio of 1:10. qPCR was performed with 2µL of cDNA using the TaqMan™ Fast Advanced Master Mix (Thermo Fisher Scientific) according to manufacturers’ instructions with the following TaqMan probes human VEGFA (Hs00900055_m1; Thermo Fisher Scientific), mouse VEGFA (Mm00437306_m1; Thermo Fisher Scientific), human GAPDH (Hs02786624_g1; Thermo Fisher Scientific), mouse GAPDH (Mm99999915_g1, Thermo Fisher Scientific). Mouse eyeballs were enucleated and stored in 4°C in RNALater^®^ until dissection. Retina was dissected from injected animals and placed in lysis buffer and sonicated for 2 seconds prior to RNA extraction and qPCR as previously described.

### Western blot

At 48 hours after transfection, cells were harvested and collected in Pierce^™^ RIPA buffer (Thermo Fisher Scientific) containing Pierce™ Phosphatase Inhibitor Mini Tablets (Thermo Fisher Scientific). Cell lysate was sonicated for 5 seconds and centrifuged for 15 minutes at 13300rpm at 4°C. The supernatant was collected, and protein levels were quantified using the Pierce™ BCA Protein Assay Kit (Thermo Fisher Scientific) as per manufacturers’ instructions. For Western blot, 10µg of protein mixed with NuPAGE™ LDS Sample Buffer (4X) (Thermo Fisher Scientific) and loaded in NuPAGE™ 4 to 12% Mini Protein Gel (Thermo Fisher Scientific) for gel electrophoresis at 150V for 30 minutes. Transfer to Amersham™ Hybond™ PVDF membrane (Cytiva) was performed at 30V for 1 hour. Protein blot was blocked with 5% skim milk for 1 hour, followed by incubation with HA-tag (C29F4) Rabbit mAb (Cell Signaling Technology^®^) or Anti-Actin Antibody, clone C4 (Merck) overnight at 4°C. Blots were then stained with Goat anti-Mouse IgG (H+L) Cross-Adsorbed Secondary Antibody, HRP (Invitrogen™) or Goat anti-Rabbit IgG (H+L) Secondary Antibody, HRP (Invitrogen™) for 2 hours at room temperature. Blot was incubated with Pierce™ ECL Western Blotting Substrate for 1 minute before imaging using Bio-Rad ChemiDoc. All washes were performed with 1× PBST for 5 minutes, repeated 3 times.

### ELISA

At 48 hours after transfection, 100µL of supernatant was collected and ELISA was performed using the Human VEGF DuoSet ELISA (R & D Systems) as per manufacturers’ instructions. Briefly, 96-well plates were incubated with Capture Antibody in ELISA Plate coating buffer overnight at room temperature. Plates were then blocked with Reagent Diluent for 1 hour at room temperature, and 100µL of supernatant was incubated for 2 hours at room temperature. Then, Detection antibody was added and incubated for 2 hours at room temperature. After incubation with Streptavidin-HRP and Substrate reagent for 20 minutes each, the provided stop solution was added, and plate was read using the Tecan Spark^®^ Micro plate reader at 450nm with reference wavelength 570nm.

### Immunohistochemistry

Retinal flatmounts were washed with 1× PBS containing 0.5% Triton-X-100 (PBST). After washing with PBST, flatmounts were incubated in anti-rhodopsin antibody (1:500; Abcam) with 2% donkey serum and 2% PBST in 4°C for 72 hours. This was followed by incubation in Alexa Fluor 488 AffiniPure Donkey Anti-Mouse IgG (H+L) (1:500; Jackson ImmunoResearch) at room temperature for 2 hours. After washing, the flatmounts were stained with Anti-Laminin-2 (α-2 Chain) antibody, Rat monoclonal (1:250; Merck) and incubated in 4°C for 5 nights. This was followed by incubation in Alexa Fluor^®^ 647 AffiniPure Donkey Anti-Rat IgG (H+L) (1:250; Jackson ImmunoResearch) at room temperature for 2 hours. Then, the retinae were washed and incubated in mCherry Monoclonal Antibody (16D7), Alexa Fluor 594 (1:500; Thermo Fisher Scientific) at 4°C for 72 hours. Retinal cryosections were blocked using serum-free protein block (Agilent Dako) at room temperature for 1 hour, followed by incubation at 4°C overnight with primary antibody as previously described. Secondary antibody incubation was performed at room temperature for 1 hour as previously described. Slides were stained with NucBlue™ Live ReadyProbes™ Reagent (Hoechst 33342; Thermo Fisher Scientific) for 10 minutes before mounting as previously described and imaged on the Stellaris 5 confocal microscope.

### hESC Maintenance and Retinal Differentiation Culture

hESCs were maintained on feeder-free conditions on E8 (Thermo Fisher) and Geltrex-coated six-well plates. In brief, for retinal differentiation hESCs were maintained until 90–95% confluent, then FGF-free medium was added to the cultures for 2 days followed by a neural induction period where RPE and retinal vesicles appeared. NRVs were excised and kept in low-binding 96-well plates for maturation.

### AAV transduction of retinal organoids

Retinal organoids were maintained in ALT90 medium as previously described (West et al., 2022) and AAV transduction was performed during the 15-19 weeks of differentiation. On the day of transduction, AAV vectors were diluted in 300µL of fresh ALT90 medium at titer of 1 × 10^11^ vg/organoids. The following day treated wells were supplemented with 700µL of fresh ALT90 medium. Transduced organoids were incubated for 72 hours in the AAV vector-containing medium. After the 72-hour incubation, the vector-containing medium was aspirated, and 1mL of fresh ALT90 medium was added to each well. Transduced organoids underwent 6 weeks culture until harvesting for experimental procedures.

### Immunohistochemistry of retinal organoids

Retinal organoids were fixed for 45 minutes in 4% paraformaldehyde and incubated overnight in 20% sucrose, prior to embedding in OCT. Cryosections (14 µm thick) were collected for analysis and preserved at - 20°C. Cryosections were blocked in 5% goat serum and 1% bovine serum albumin in PBS for 2 hours. Primary antibodies (ARR3: 1:200, Rhodopsin: 1:1000, VSX2: 1:200, CRALBP: 1:200, PROX1:1:200, and HuCHuD:1:500) were incubated overnight at 4°C in blocking solution with 5% goat or donkey serum. After washing of primary antibody with PBS sections were incubated with secondary antibody in blocking solution for 2 hours at RT, washed and counter-stained with DAPI (Sigma-Aldrich). Alexa fluor 488 and 546 secondary antibodies (Thermo Fisher Scientific) were used at a 1:500 dilution.

### Intravitreal injections

All animal procedures were approved by the Animal Ethics Committees of University of Melbourne and conducted in accordance with The Association for Research in Vision and Ophthalmology (ARVO) Statement for the Use of Animal in Ophthalmic and Vision Research. Ethics approval was obtained from The Florey Institute of Neuroscience and Mental Health Animal Ethics Committee (22-036-UM). The Akimba mice developed by Rakoczy *et al*. (46) and the heterozygous Kimba mice (*hVEGFA*^+/-^) were used for this study. Mice were bred in the Melbourne Brain Centre animal facility (Kenneth Myer Building, Parkville, VIC, Australia), and were housed in a well-ventilated environment that was kept at a constant room temperature of 21°C, with a 12-hour diurnal light-dark cycle (lights on: 7am, off: 7pm; room illumination < 50 lux). Mouse chow (Barastoc, Melbourne, VIC, Australia) and water were provided ad libitum. 6-week-old Kimba mice were anesthetized with ketamine (80mg/kg) and xylazine (10mg/kg) solution mix (Troy Laboratory, Smithfield, NSW, Australia; i.p. with 30G needle) and given a drop of 0.5% proxymethacaine hydrochloride (Alcaine™, Alcon Laboratories, Frenchs Forest, NSW, Australia) and 0.5% tropicamide (Mydriacyl™, Alcon Laboratories). Baseline fundus fluoresceine angiography (FFA) were taken (as described below), and animals intravitreally injected with 1µL of AAV2.7m8 (1 × 10^10^ vg/µL) viruses using Hamilton syringe attached to a 33-gauge dental needle. Post injection, mice were subcutaneously administered 0.3mL of atipamezole hydrochloride (0.25mg/mL) and allowed to recover on heating pads with food and water *ad libitum*.

### Fundus fluorescein angiography and optical coherence tomography

Retinal vasculature was assayed at baseline prior to intravitreal injection of AAVs and 8 weeks after AAV-treatment using fundus fluorescein angiography (FFA) (Spectralis, Heidelberg Engineering, Heidelberg, Germany) immediately after intraperitoneal injection of sodium fluorescein (1%, 100 µL/kg, fluorescite 10%; Alcon Laboratories). Five images (central, nasal superior, temporal superior, nasal inferior, and temporal inferior) were obtained for each eye and analysed using AngioTool(47) for vessel density and branch points using default settings.

*In vivo* retinal structure was assayed with spectral domain optical coherence tomography (SD-OCT) (Spectralis^®^, Heidelberg Engineering, Heidelberg, Germany) with general anaesthesia, mydriasis and corneal hydration (Systane, Alcon Laboratories) maintained. During image acquisition, retinal volumes (8.1 × 8.1 × 1.9 mm) centred on the optic nerve head were acquired each consisting of 121 evenly spaced horizontal B-scans. Each B-scan comprised of 768 A-scans with 3.87 µm axial depth resolution, 9.8 µm lateral resolution. OCT scans were captured with an automated real-time tracking average of 6 frames, at an average speed of 85,000 A-scans per second. The Heidelberg Eye Explorer 2 OCT reader plugin (Heidelberg Engineering) was used for analysis. All retinal layers (retinal nerve fibre layer, RNFL; ganglion cell inner plexiform layer, GCIPL; inner nuclear layer, INL; outer plexiform layer, OPL; outer nuclear layer, ONL; photoreceptor segments, PR segment) as well as total retinal thickness (TRT) were automatically segmented (Heidelberg Eye Explorer 2 software). A circular ring around the optic nerve head (Early Treatment Diabetic Retinopathy Study [ETDRS] outer ring) was employed for analysis.

### Determination of candidate off-target genes

Off-targets of the gRNAs were predicted using a sequence-based approach. gRNAs in length of 30 bps were first aligned to the human transcriptome (Grch38 cdna from emsembl release 100) using blastn (megablast). Flexible options were set to capture the potential offtargets which could tolerate up to 7 mismatches in this step, i.e., max_target_seqs = 10000, evalue = 10000, word_size = 5, perc_identity = 0.6. Secondly, the candidates were further filtered with the threshold that at least 13 bp perfect match should exist in each alignment.

### Bulk RNA-seq data analysis

Raw data processing was performed using FASTQC (48) to check the reads quality and Trimmomatic (49) tool to further remove the bases and reads with low quality, adaptors and barcodes. The clean reads were aligned to the reference human genome (version GRCh38), utilizing STAR v2.7 (50) as the aligner. For RNA quantification, RSEM (51) was employed to normalize raw read counts, transforming them into TPM (transcripts per million) values, providing a relative expression level that is theoretically comparable across samples. The magnitude (log2-transformed fold change) and significance (P-value) of differential expression between groups were then calculated. Differentially expressed genes (DEGs) were selected based on the criteria of a p-value < 0.05 and |log2 fold change| ≥ 1. Gene set enrichment analysis (GSEA) (52), focused on Gene Ontology(53), was conducted using the R 4.3.1 Bioconductor package clusterProfiler(54) to elucidate whole transcriptomic patterns between groups. To account for multiple testing, P-values underwent adjustment through the Benjamini-Hochberg false discovery rate (BH-FDR).

### Single-cell RNA-seq data analysis

After checking the reads quality, the sequencing files were aligned to reference human genome (version GRCh38) using 10X Genomics CellRanger software (v7.1.0) count pipeline to remove non-cellular barcodes and keep the cellular barcodes and gene matrix. To further distinguish the Cas13bt3-valid and -invalid cells, FASTX-toolkit (55) was used to keep the reads with cellular barcodes and convert FASTAQ file into FASTA file. Then, Blastn (56) was used to compute the similarity between cellular reads and Cas13bt3 reference sequence.

Based on the quality of the sequencing reads, appropriate cutoff values for both the percentage of identical positions and the alignment length were employed to delineate cells as either Cas13bt3-valid or Cas13bt3-invalid. For either Cas13bt3-valid or -invalid cells, we further used Seurat v4.3 for data pre-processing, data normalization, feature selection, data scaling, dimensional reduction and clustering. To refine the characterization of cell types within each cluster, ScType (57) was employed, leveraging genesets specific to eight retinal cell types—amacrine, bipolar, cones, horizontal, macroglial, microglial, rods, and RPE— validated by Kim et al. (27). The magnitude (log2-transformed fold change) and significance (P-value) of differential expression between cell types were then calculated. Differentially expressed genes (DEGs) were selected based on the criteria of a p-value < 0.05 and |log2 fold change| ≥ 1. Gene set enrichment analysis (GSEA) (52), focused on Gene Ontology (53), was conducted using the R 4.3.1 Bioconductor package clusterProfiler (54) to elucidate whole transcriptomic patterns between groups. To account for multiple testing, P-values underwent adjustment through the Benjamini-Hochberg false discovery rate (BH-FDR).

### Statistical analysis

Statistical analyses were performed using GraphPad Prism 10. Analytical methods are described in respective figure legends with p-value indicators. For ex vivo studies, to assess the strength of the relationship between gene expression values and the number of Cas13bt3-valid read counts following log_2_ transformation in each cell type, Spearman’s rank correlation coefficient was employed. The associated *p-value* serves as an indicator of the likelihood or probability that any observed correlation is merely a result of chance.

## Acknowledgments

The authors would like to thank Liuhui Huang, Jesse Gardner-Russell and Yujie Wang for their assistance with retinal dissections, Prof Alex Hewitt and Dr Leilei Tu for providing resources for vector construction. The authors also extend their gratitude to the Vector and Genome Engineering Facility (VGEF) and the Single Cell Analytics Facility at the Children’s Medical Research Institute (CMRI) for producing AAV vectors and the scRNA-seq experiments respectively and Azenta Life Sciences for conducting RNA sequencing. Schematic in figures were designed using Biorender.com.

## Supplementary Materials

**Figure S1.**
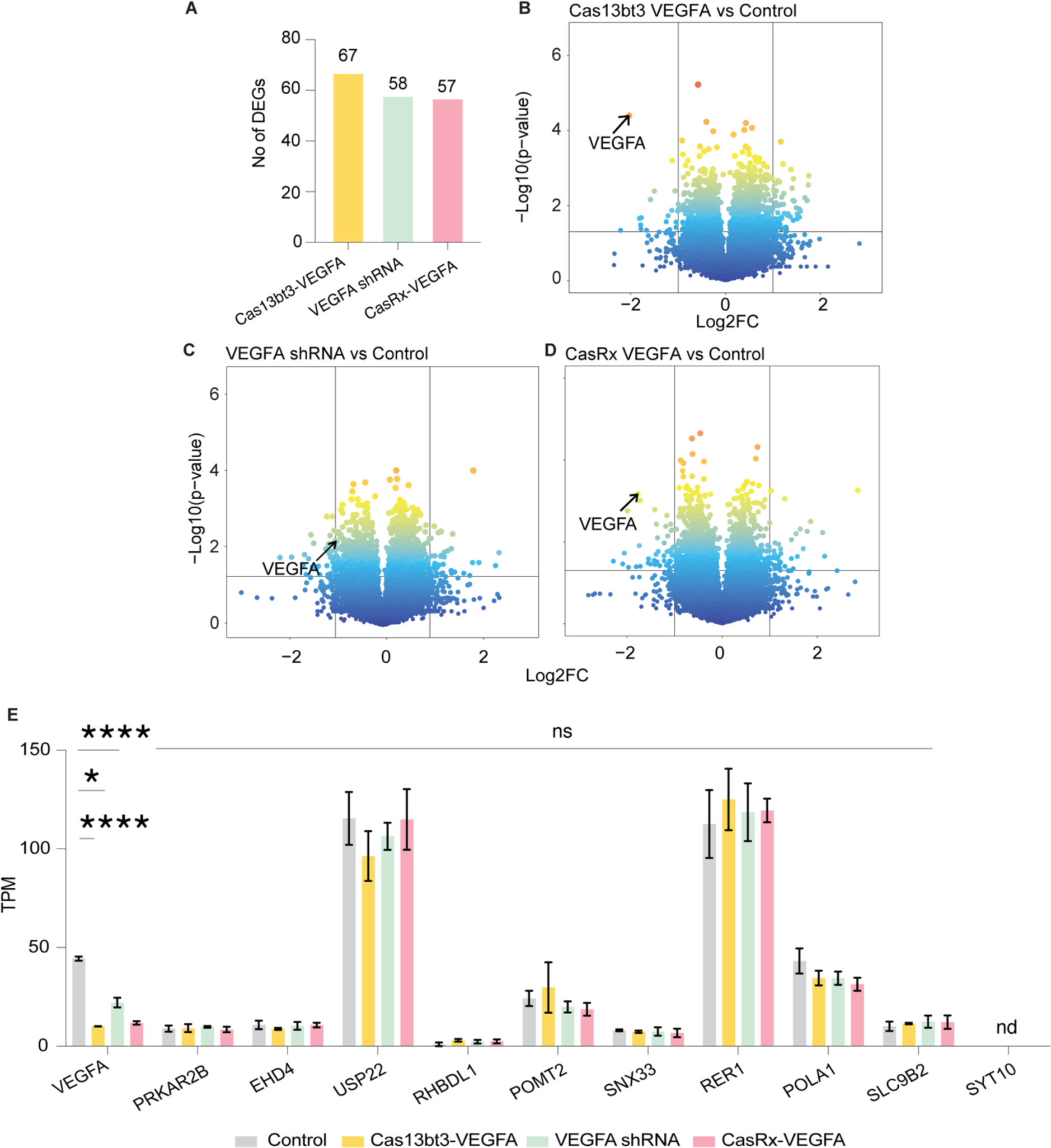
HEK293FT DEG and off-target analysis. **(A)** Number of differentially expressed genes from the VEGFA-targeting shRNA, CasRx and Cas13bt3 transfections in HEK293FT cells compared to vector control. **(B)** Volcano plots of differentially expressed genes from Cas13bt3 VEGFA sgRNA **(C)** VEGFA shRNA and **(D)** CasRx VEGFA sgRNA compared to vector control. **(E)** Plot of TPM values for VEGFA and candidate off-target genes from Cas13bt3-NT sgRNA and Cas13bt3 VEGFA sgRNA groups compared to shRNA and CasRx targeting vectors. Statistical analysis was conducted by Welch’s t test **(E)**. Data presented as mean ± SEM. ns: not significant. *p≤0.05, ****p≤0.0001.

**Figure S2.**
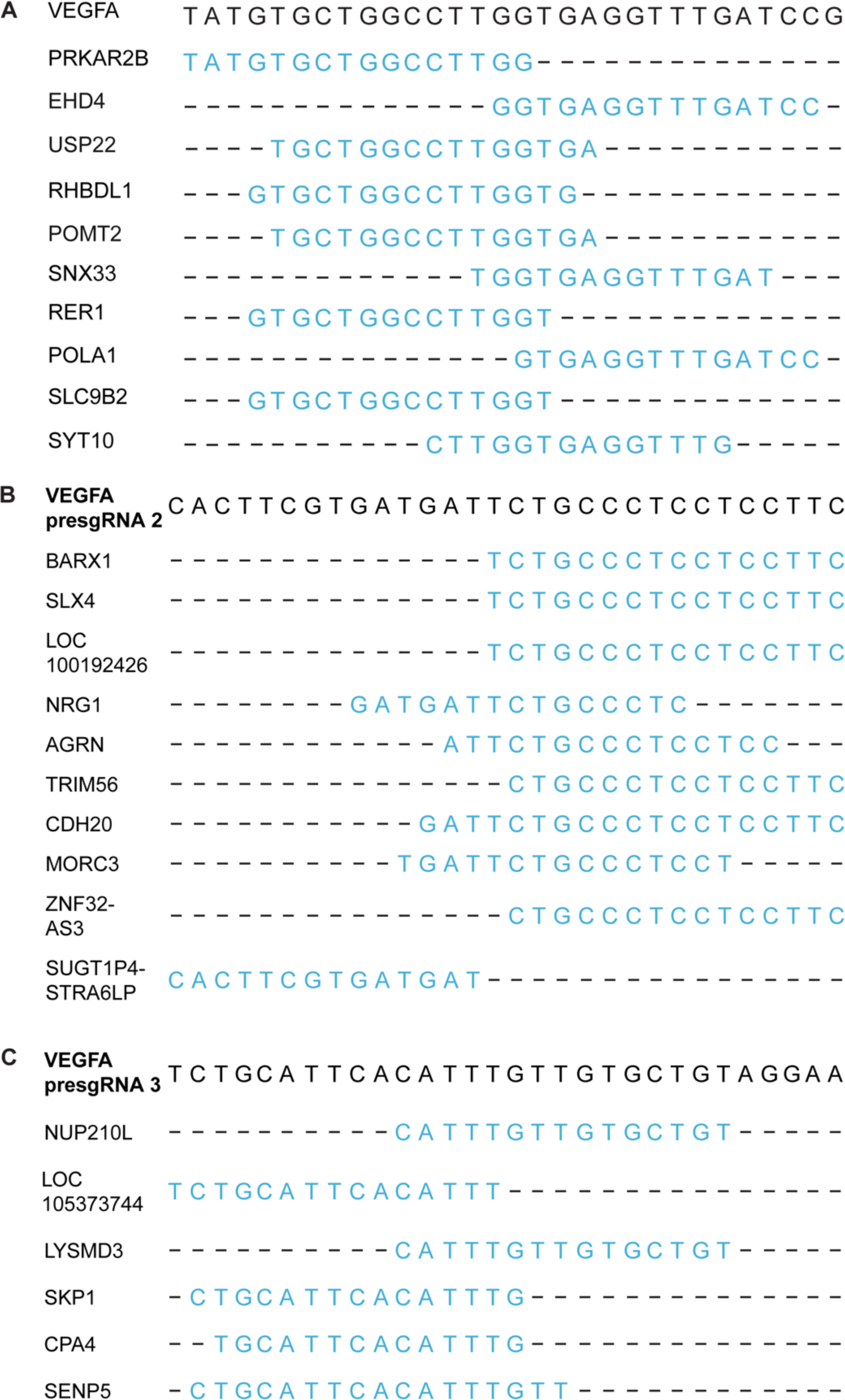
Multiplexed VEGFA knockdown using Cas13bt3. **(A)** Top off-target genes for VEGFA sgRNA 1, **(B)** VEGFA sgRNA 2 and **(C)** VEGFA sgRNA 3.

**Figure S3.**
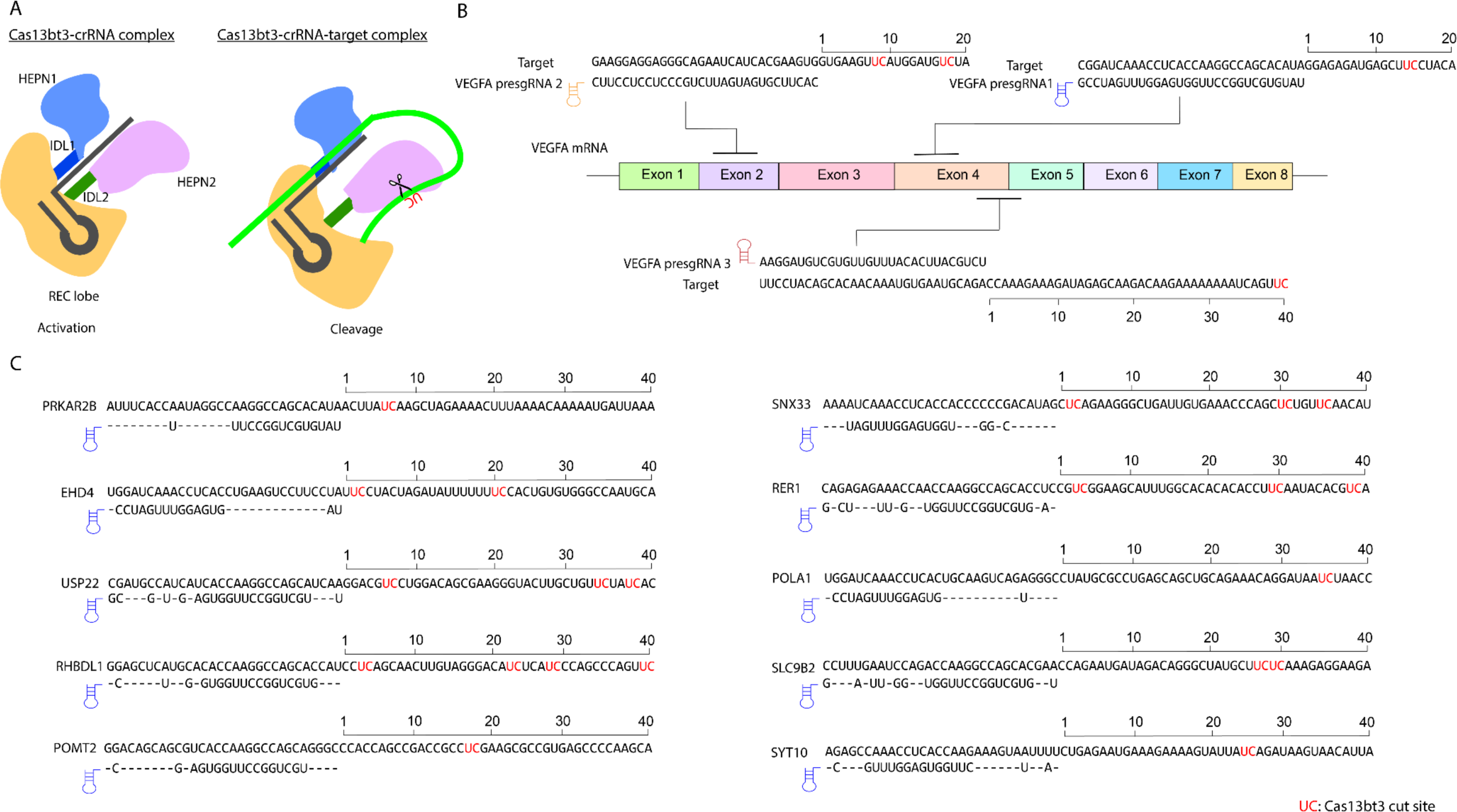
Prediction of Cas13bt3 cut sites. **(A)** Schematic of Cas13bt3-gRNA and Cas13bt3-gRNA-target complex. **(B)** Potential Cas13bt3 cut sites on VEGFA mRNA target for VEGFA sgRNA1, 2 and 3. **(C)** Potential Cas13bt3 cut sites on top 10 candidate off-target genes for VEFGA sgRNA 1.

**Figure S4.**
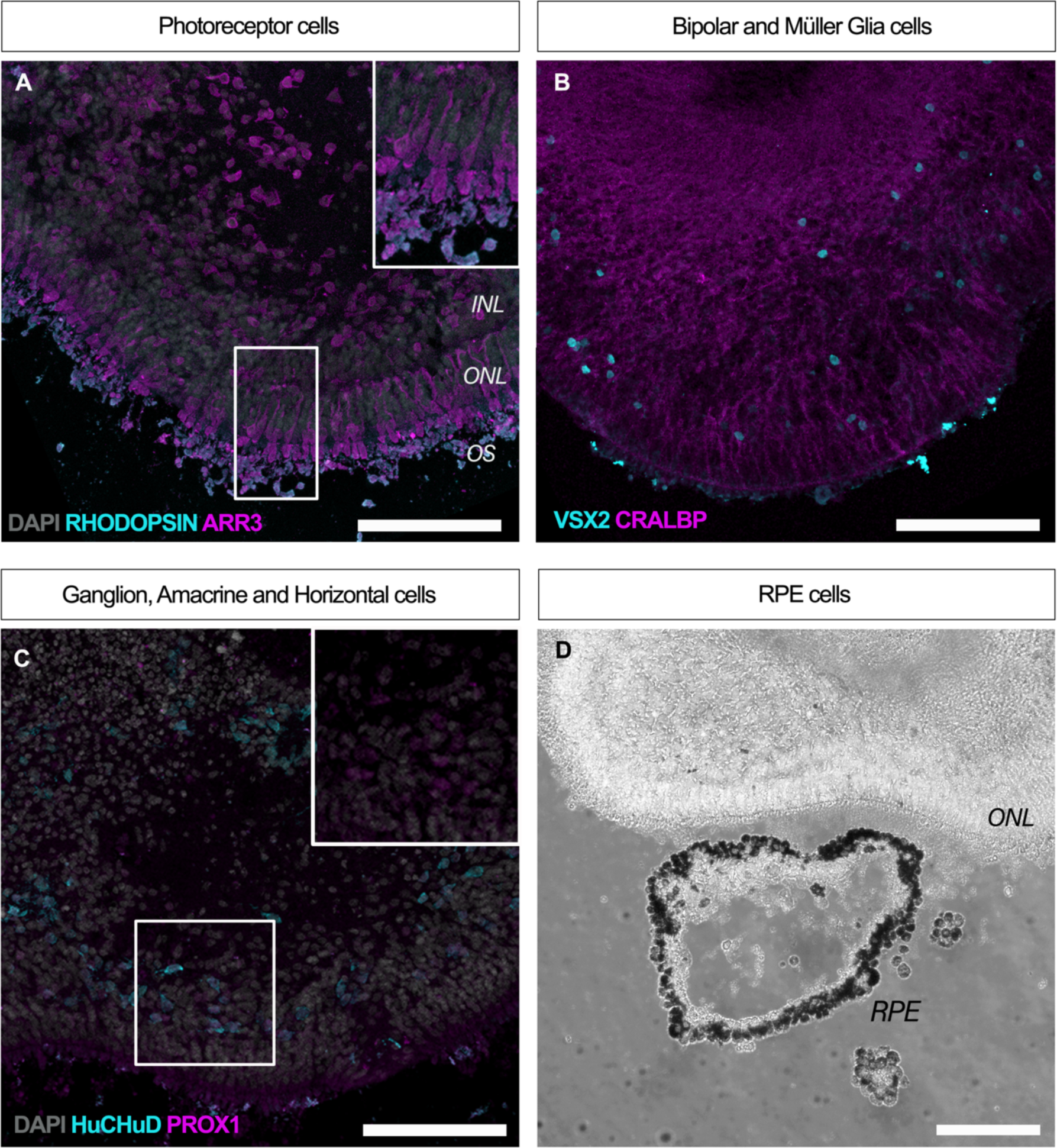
Characterization of 30 weeks retinal organoids. **(A)** Retinal organoids showed expression of mature photoreceptor markers Arrestin-3 (magenta) specific to the cone cells and Rhodopsin (cyan) localized to the outer segments (OS) of rod photoreceptor cells. **(B, C)** In the inner nuclear layer (INL) bipolar cells stained with VSX2 (cyan) and Müller glia stained with CRALBP (magenta) **(B)**, Amacrine and horizontal cells stained with PROX1 (magenta) and ganglion cells stained with HuCHuD (cyan) **(C)**. **(D)** Brightfield image of retinal organoid section showing the outer nuclear layer (ONL) and a pigmented RPE island next to it. Scale bar: 116μm. Abbreviations: INL: inner nuclear layer ONL: outer Nuclear Layer, OS: outer segment, RPE: retinal pigment epithelium.

**Figure S5.**
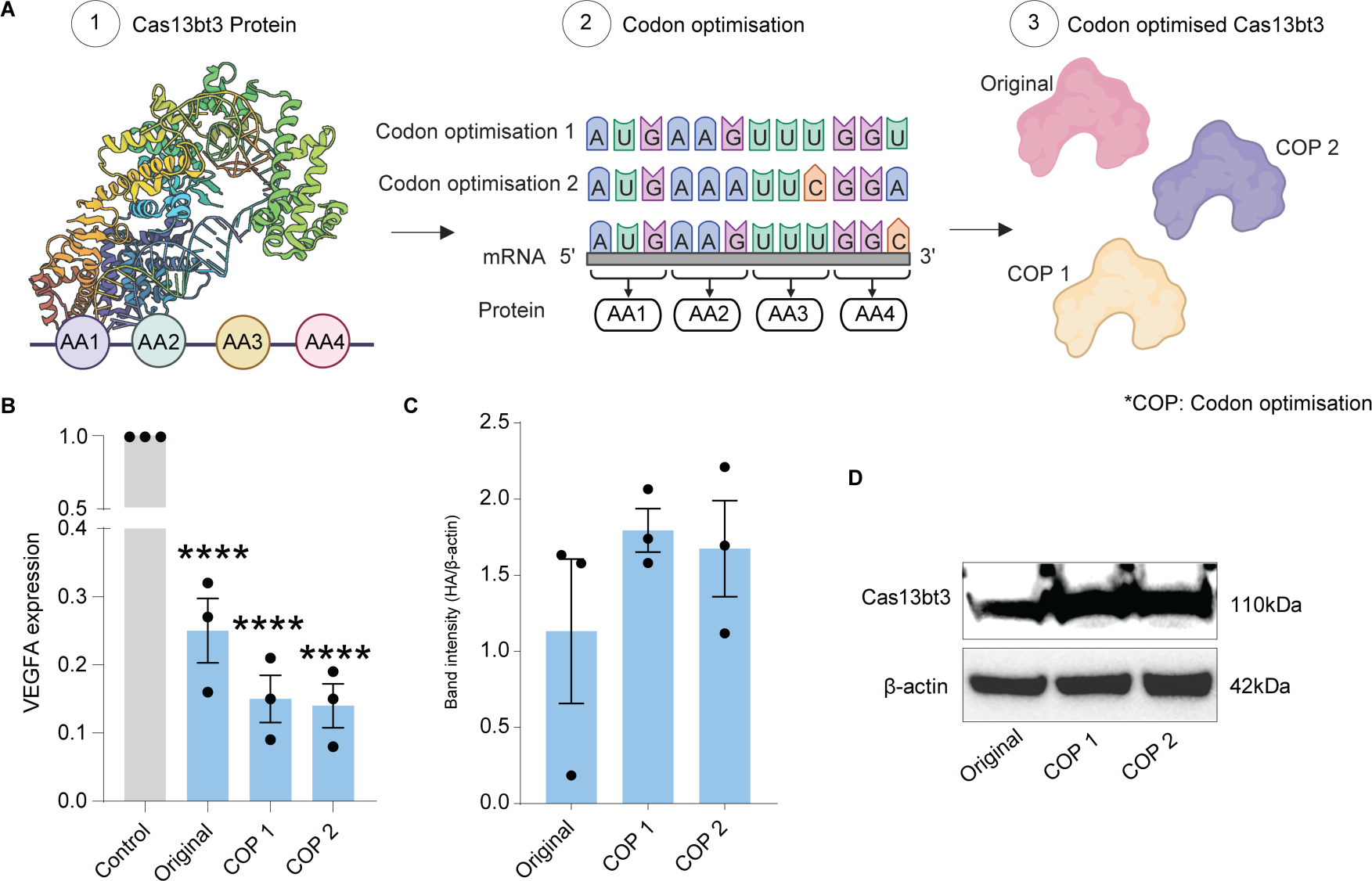
Codon optimisation of Cas13bt3. **(A)** Schematic of codon optimisation procedure from Cas13e protein sequence. **(B)** VEGFA mRNA knockdown from original and codon optimised Cas13e compared to control transfection. **(C and D)** Expression of original and codon optimised Cas13bt3 variants determined from Western blot. Statistical analysis was conducted using One-way ANOVA with Tukey’s multiple comparisons test (**B and C**). Data presented as mean ± SEM. *p≤0.05, **p≤0.01, ****p≤0.0001.

**Figure S6.**
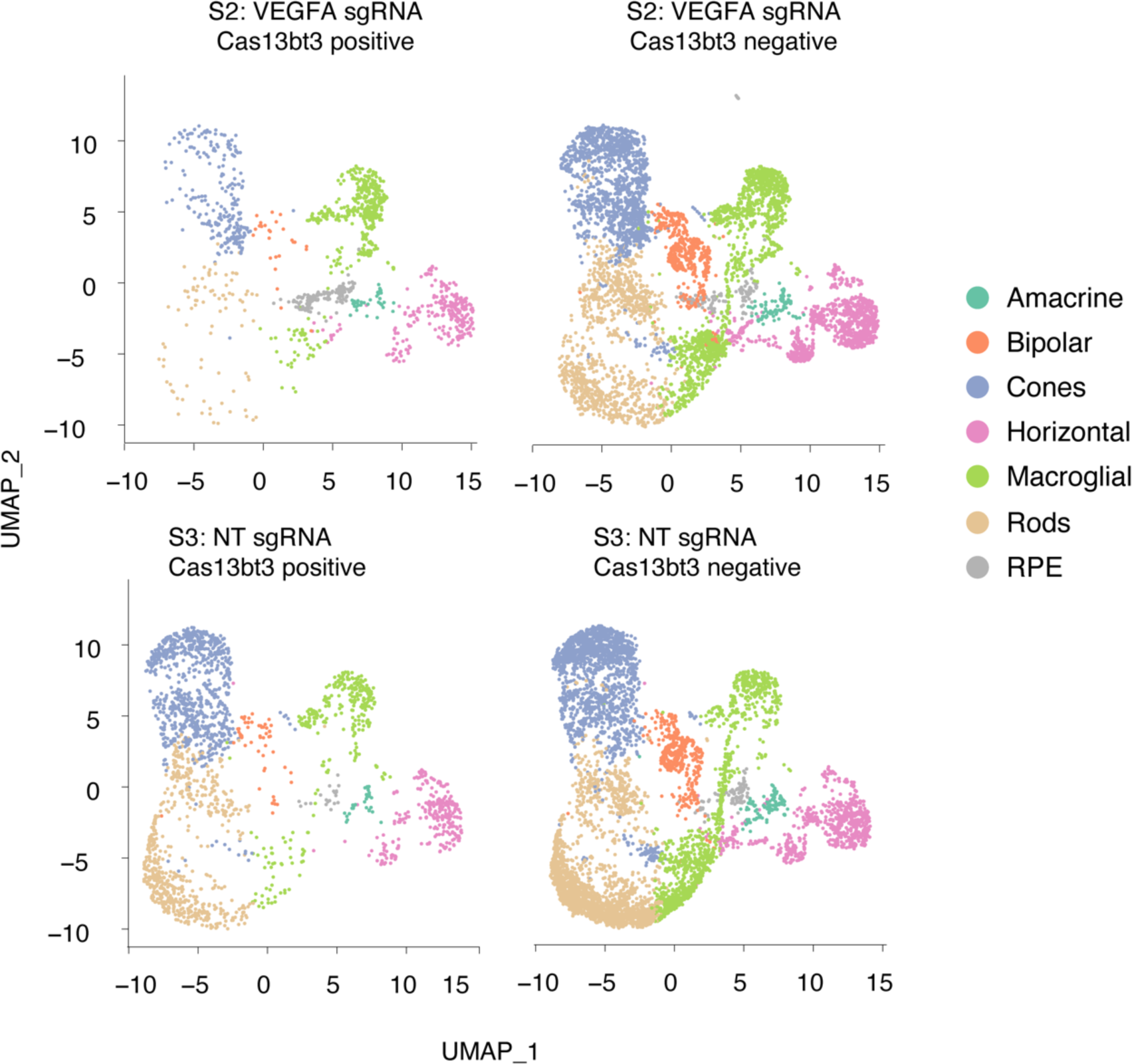
Clustering of retinal cells in retinal organoids. UMAP plots show retinal cell type clusters for the NT sgRNA and VEGFA sgRNA treatment groups highlighting Cas13bt3 positive and negative cells.

**Figure S7.**
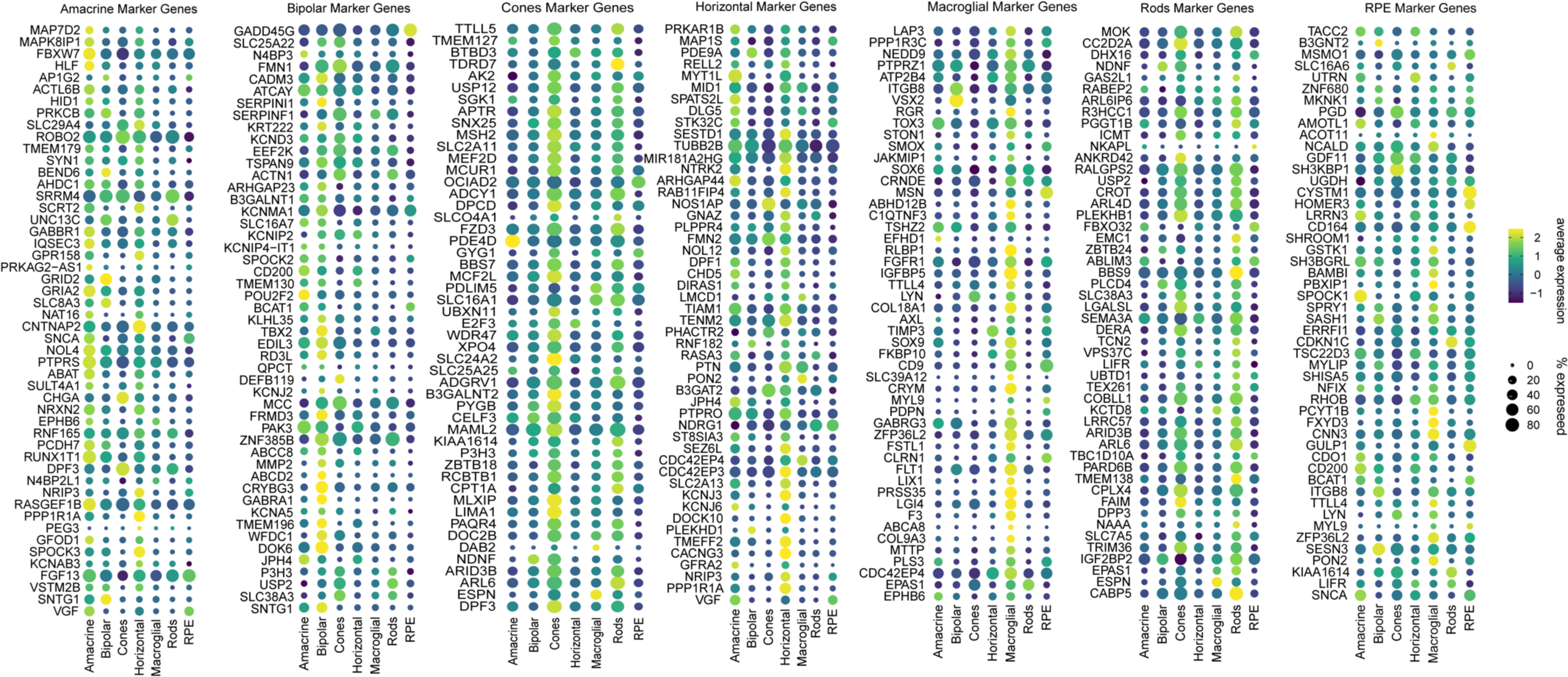
Dot plots of retinal cell marker genes for respective cell types in 3D retinal organoids. Complete list of cell marker genes can obtained from Kim et al., 2023 (*27*).

**Figure S8.**
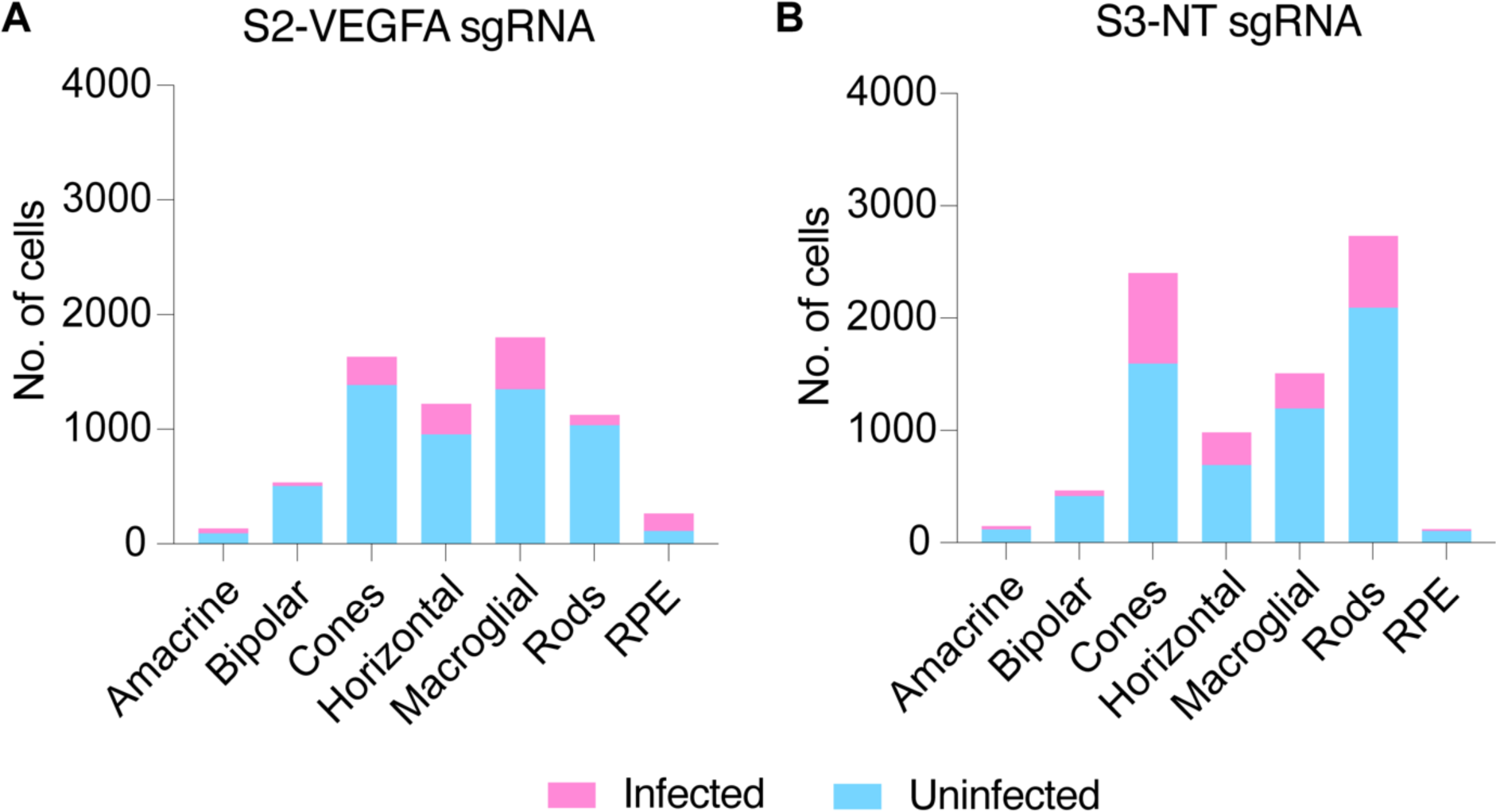
Infection rate of retinal organoids. **(A)** Number of retinal cells infected and uninfected for the S2-VEGFA sgRNA group and **(B)** S3-NT sgRNA group.

**Figure S9.**
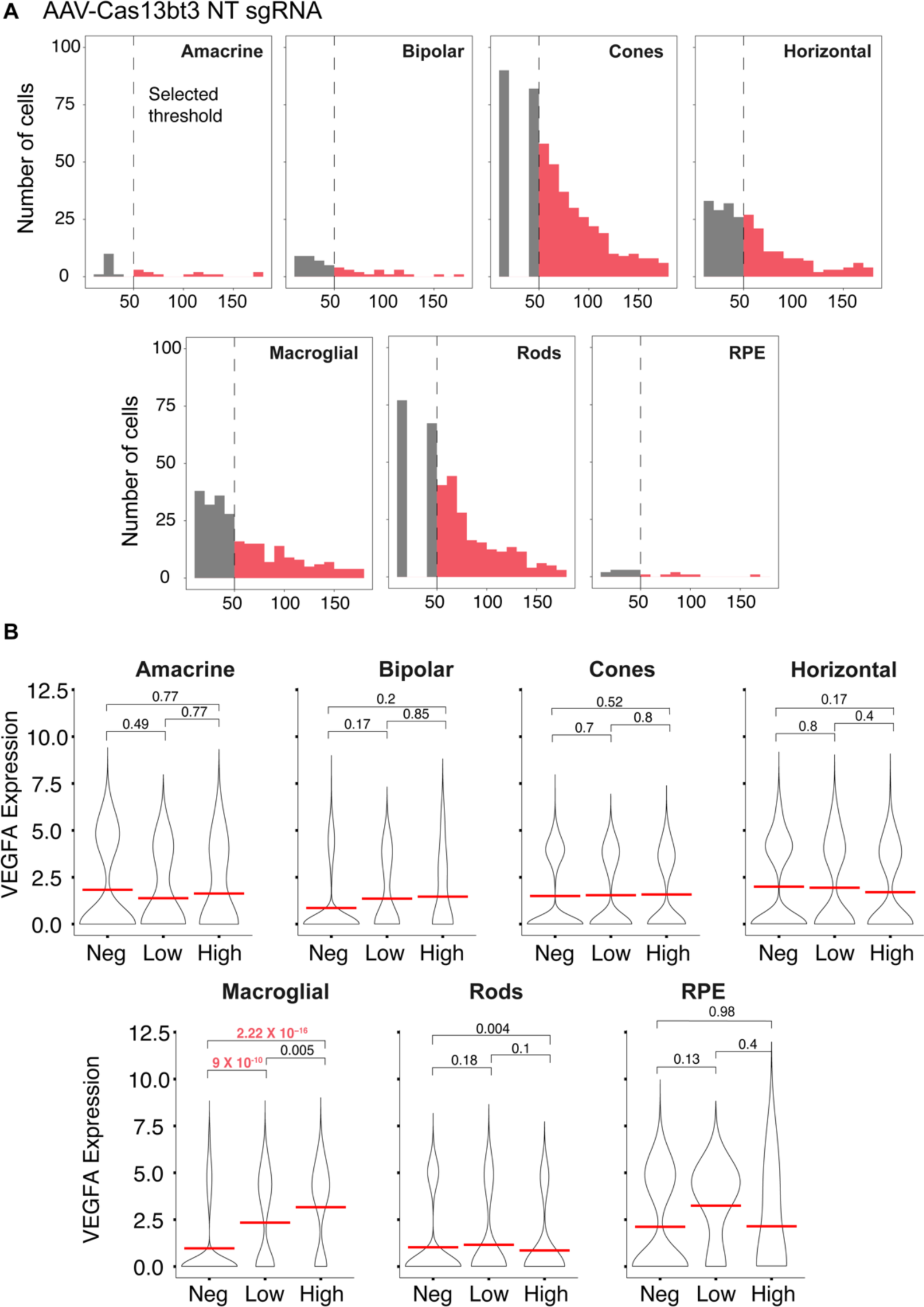
Cas13bt3-NT sgRNA treatment in retinal organoids. **(A)** Expression of Cas13bt3 by number of reads across retinal cell types in NT sgRNA treatment group. **(B)** Violin plots of VEGFA expression in cells with low and high expression of Cas13bt3 against Cas13bt3 negative cells showed no significant differences.

**Figure S10.**
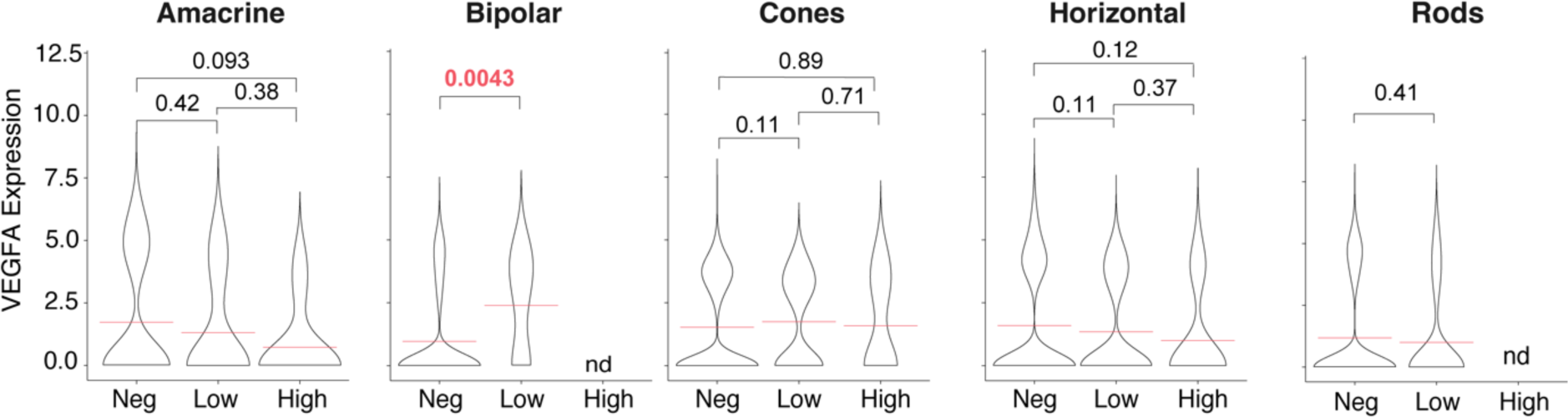
VEGFA expression in retinal cells with different levels of Cas13bt3 expression. Violin plots of VEGFA expression against the level of Cas13bt3 expression across retinal cell types compared to Cas13bt3 negative cells.

**Figure S11.**
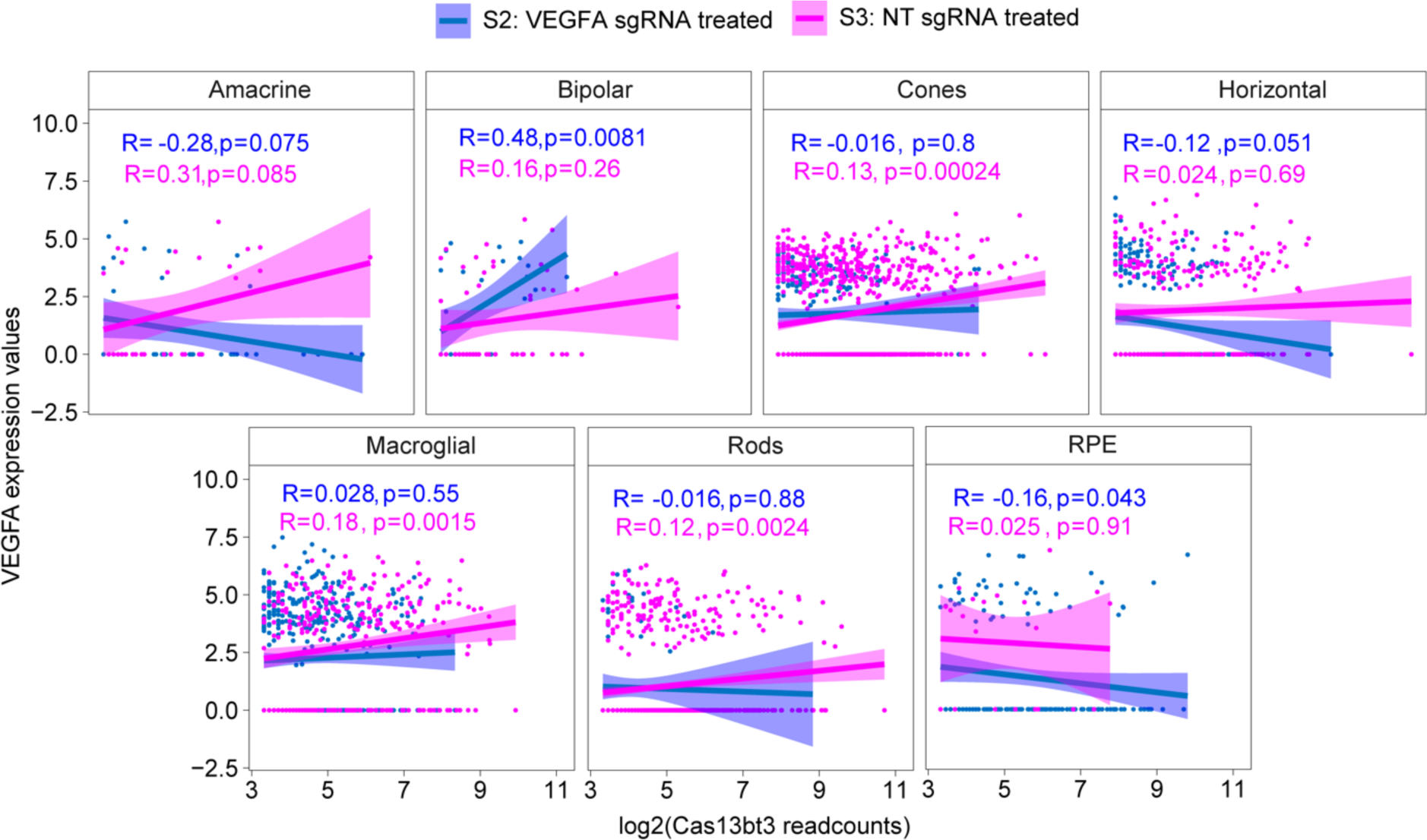
Correlation of Cas13bt3 expression with VEGFA expression. The expression of Cas13bt3 was plotted against the expression of VEGFA mRNA across cell types in NT sgRNA and VEGFA sgRNA groups.

**Figure S12.**
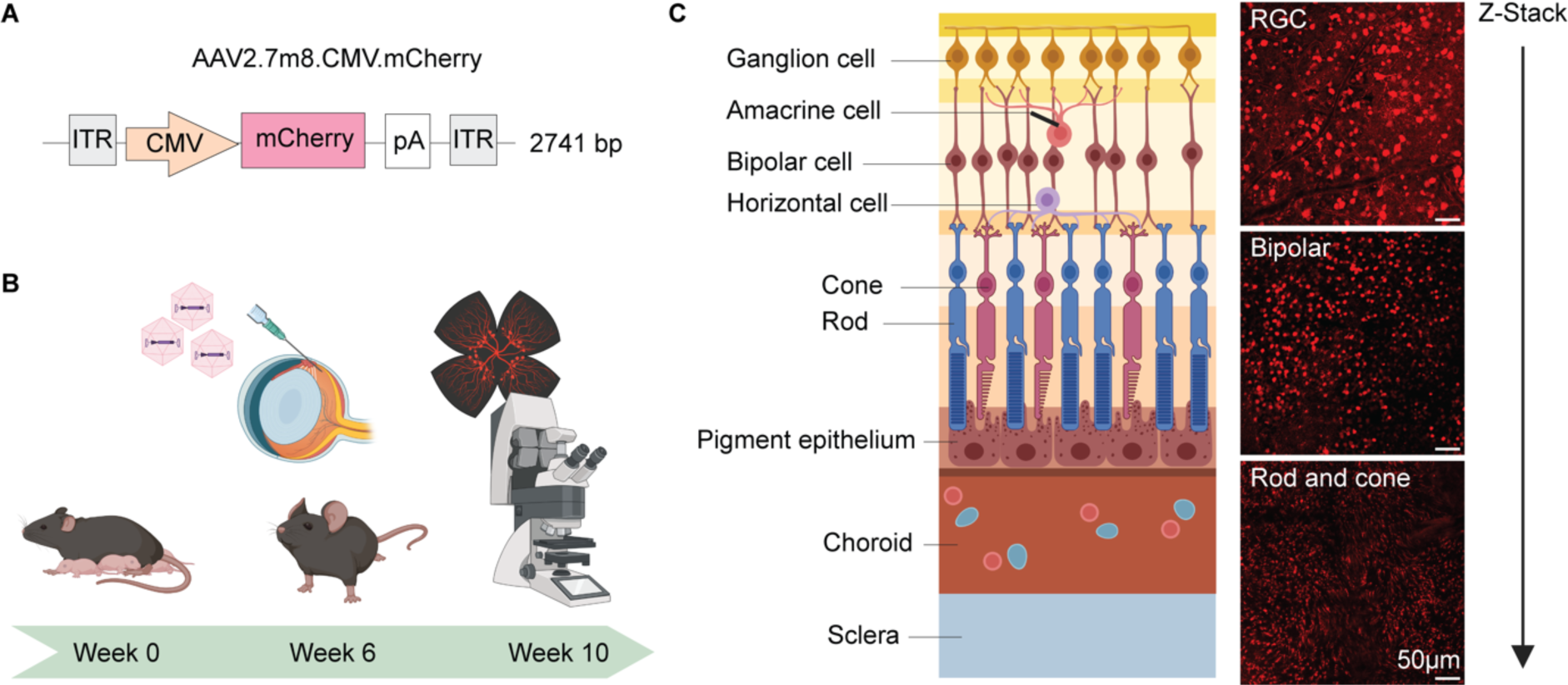
Intravitreal delivery of AAV2.7m8 achieves strong transduction of mice photoreceptors. **(A)** Schematic of AAV2.7m8 virus carrying mCherry reporter. **(B)** Schematic of experimental procedure for intravitreal injection of C57J/BL6 mice eyes. **(C)** Confocal microscope images of retinal flatmount 4 weeks post intravitreal injection, showing transduction of ganglion, bipolar, cone and rod cells with a reference image of retinal cell layers. RGC: retinal ganglion cell.

**Figure S13.**
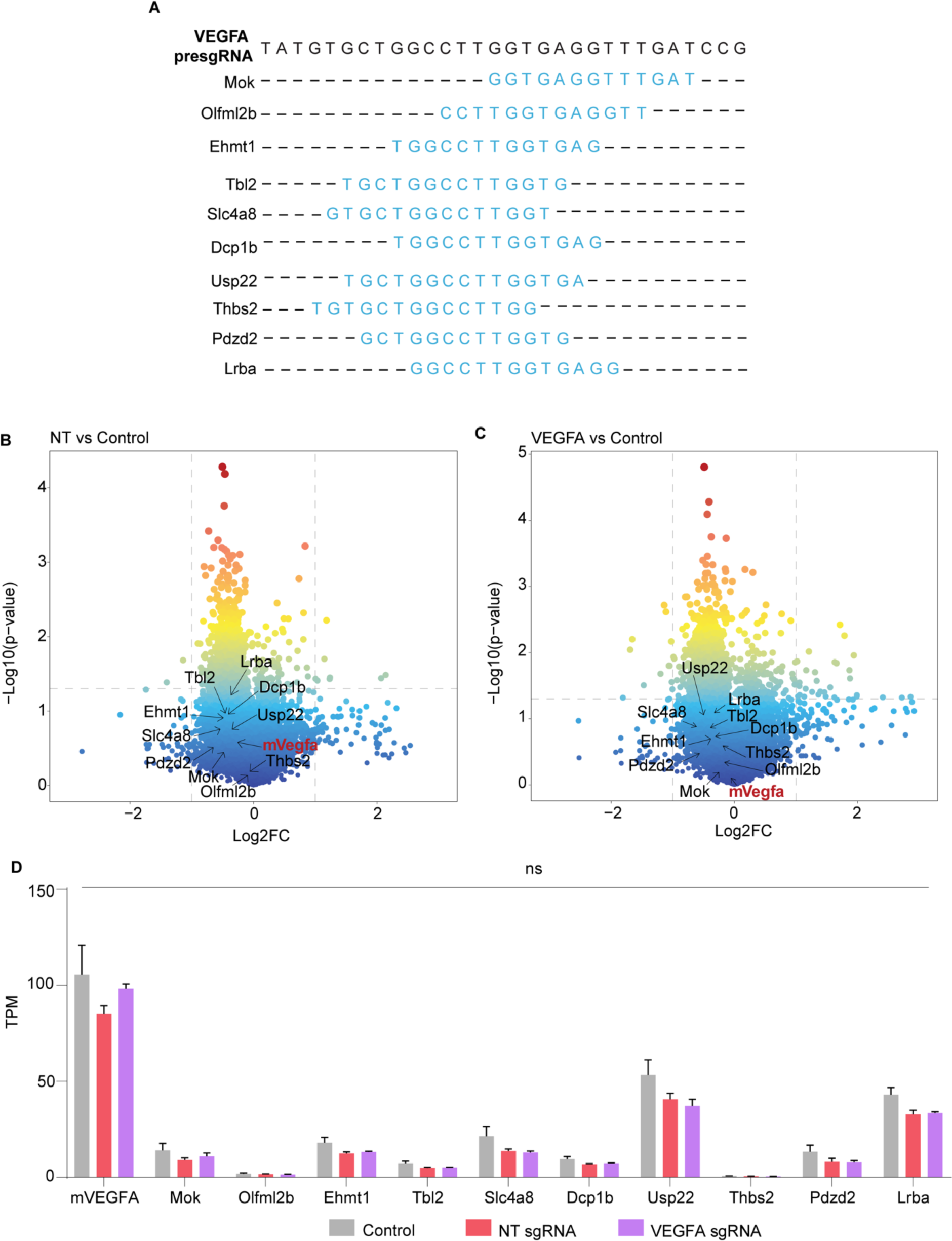
Cas13bt3 off-target effects in Kimba mice. **(A)** Schematic of candidate off-target genes to VEGFA sgRNA in mouse transcriptome. Volcano plots of differentially expressed genes from Kimba mice retinas treated with AAV2.7m8 carrying **(B)** Cas13bt3-NTsgRNA and **(C)** Cas13bt3-VEGFAsgRNA compared to vector controls. **(D)** Expression in TPM of mouse VEGFA and candidate off-target genes from RNA-seq.

**Figure S14.**
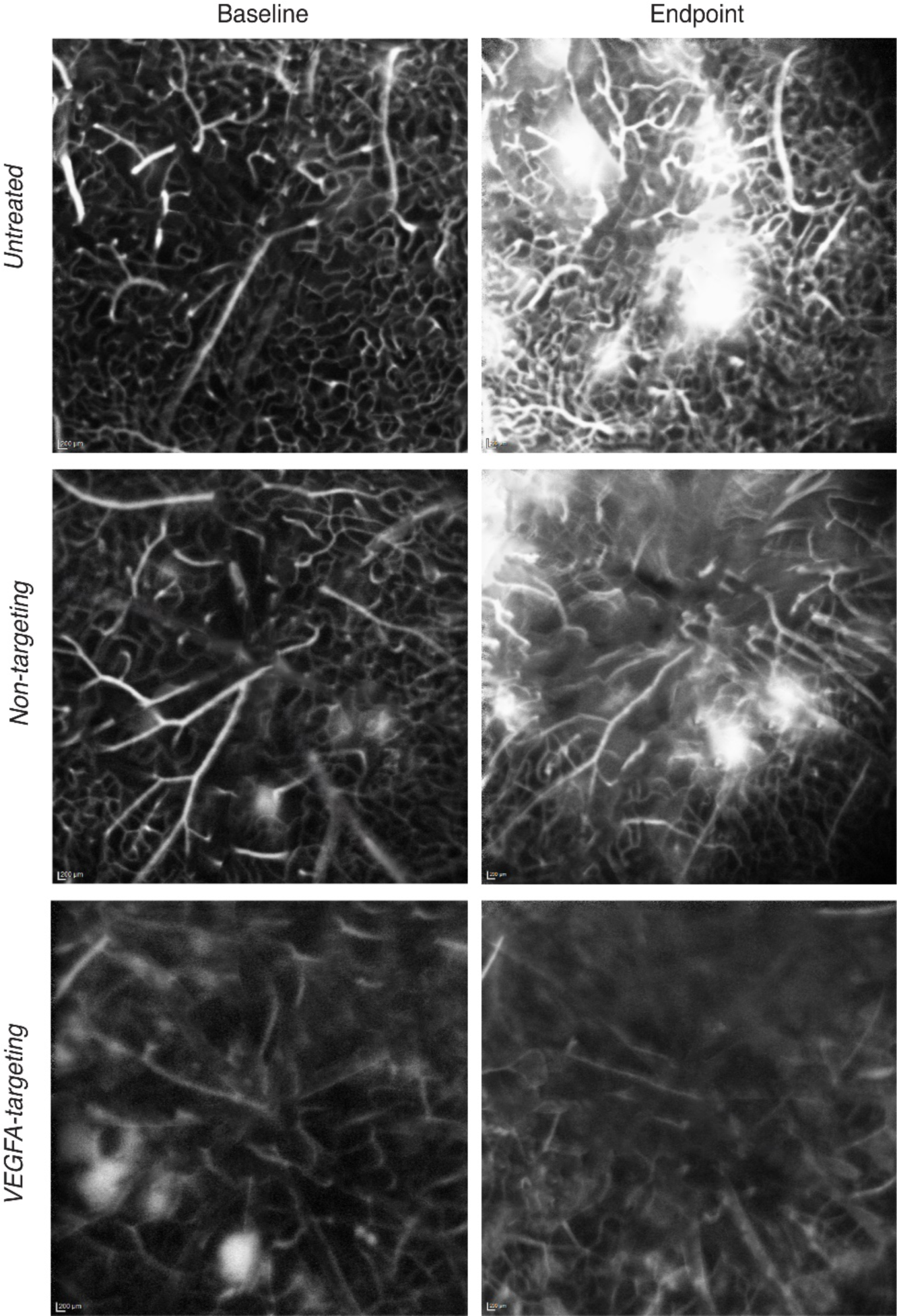
Original fluorescein angiography (FFA) images from Kimba mice retinae. Representative images of FFA collected from Kimba mice at baseline and endpoint and used for AngioTool analyses.

**Figure S15.**
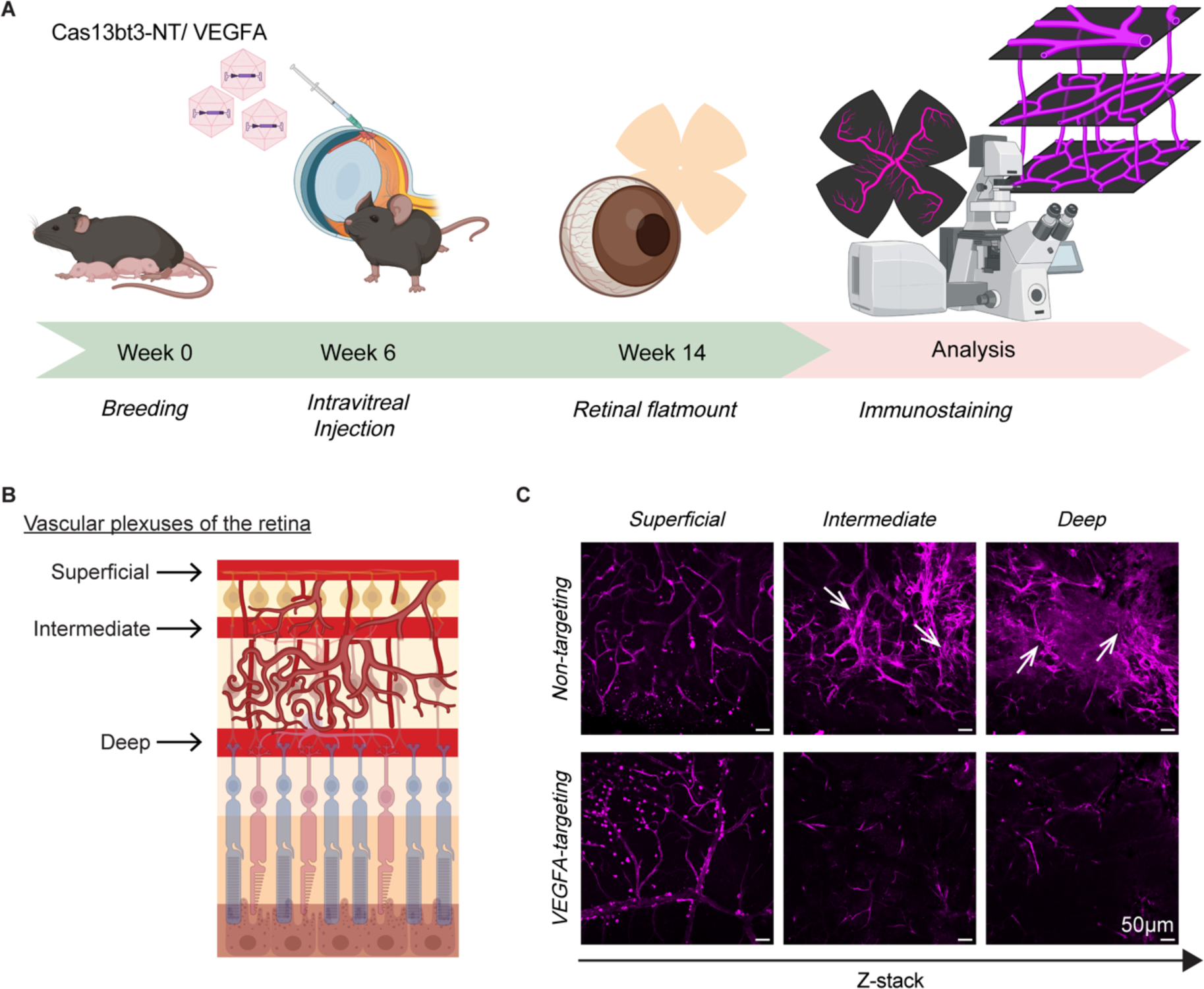
Imaging of retinal vascular plexuses Kimba mice treated with Cas13bt3-NTsgRNA and Cas13bt3-VEGFAsgRNA. **(A)** Schematic of experimental procedure for imaging retinal vascular plexuses. **(B)** Schematic of the different retinal plexuses. **(C)** Confocal imaging of the retinal plexuses, 8 weeks after Cas13bt3 treatment, immunolabelled with laminin. Magnification 40X, scale bar: 50µm. White arrows indicate leaky vessel structure.

**Figure S16.**
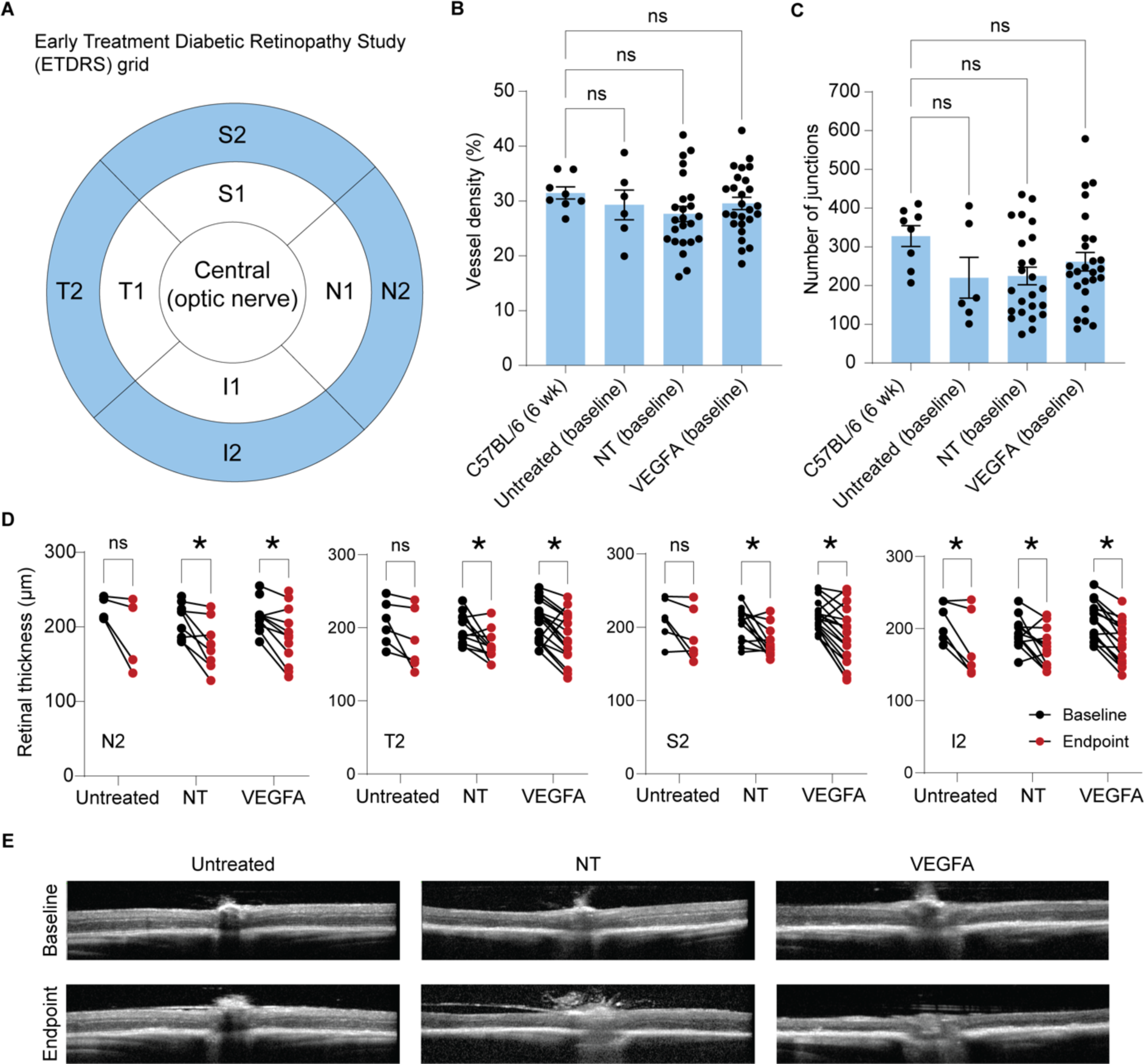
Retinal thickness readings from optical coherence tomography (OCT) in Kimba mice. **(A)** Early Treatment of Diabetic Retinopathy Study (ETDRS) grid showing segments of the retina analysed using OCT. C: Central (optic nerve); N1: Inner nasal; N2: Outer Nasal; T1: Inner temporal; T2: Outer temporal; S1: inner superior; S2: Outer superior; I1: Inner inferior; I2: Outer inferior. **(B)** Vessel density and **(C)** number of junction readings from FFA images of Kimba mice at baseline compared to wildtype (C57BL/6) mice. **(D)** Plot of difference in total retinal thickness in untreated and treated Kimba mice between baseline and endpoint. **(E)** Representative OCT images from untreated and treated Kimba mice at the endpoint.

